# Endothelial tight junctions and cell-matrix adhesions reciprocally control blood-brain barrier integrity

**DOI:** 10.1101/2025.09.06.673321

**Authors:** Gergő Porkoláb, Lucien Lemaitre, Beatrix Magyaródi, Imola Rajmon, Tibor Novák, Bálint H. Kovács, Ilona Gróf, Anikó Szecskó, Kinga Dóra Kovács, Enikő Farkas, Boglárka Kovács, Inna Székács, Attila G. Végh, Yosuke Hashimoto, Chris Greene, Adam McGlinchey, Maxime Culot, David C. Henshall, Kieron J. Sweeney, Donncha F. O’Brien, Szilvia Veszelka, Miklós Erdélyi, Matthew Campbell, Róbert Horváth, Mária A. Deli

**Affiliations:** Institute of Biophysics, Biological Research Centre, Hungarian Research Network, H-6726 Szeged, Hungary; Smurfit Institute of Genetics, Trinity College Dublin, D02 VF25 Dublin, Ireland; FutureNeuro Research Ireland Centre, Smurfit Institute of Genetics, School of Genetics and Microbiology, Trinity College Dublin, D02 VF25 Dublin, Ireland; Nanobiosensorics Laboratory, Institute of Technical Physics and Materials Science, Centre for Energy Research, Hungarian Research Network, H-1121 Budapest, Hungary; Doctoral School of Biology, University of Szeged, H-6720 Szeged, Hungary; Department of Optics and Quantum Electronics, University of Szeged, H-6720 Szeged, Hungary; Department of Physiology & Medical Physics, RCSI University of Medicine and Health Sciences, D02 YN77 Dublin, Ireland; FutureNeuro Research Ireland Centre, Department of Physiology & Medical Physics, RCSI University of Medicine and Health Sciences, D02 YN77 Dublin, Ireland; Laboratoire de la Barriére Hémato-Encéphalique, Université d’Artois, 62307 Lens, France; Department of Neurosurgery, Beaumont Hospital, D09 V2N0 Dublin, Ireland

## Abstract

Brain endothelial cells (ECs) rely on mechanical cues to provide a physical barrier that protects the brain. Yet how ECs integrate forces to establish and maintain the blood-brain barrier (BBB) remains poorly understood. Here, we show that the two main endothelial force-bearing systems, tight junctions and cell-matrix adhesions, reciprocally control BBB integrity. Using a combination of super-resolution imaging and biophysical techniques, we reveal increasing mechanical loads on cell-cell junctions vs. cell-matrix adhesions in human stem cell-derived ECs during BBB maturation. This force redistribution is enabled by cytoskeletal remodeling, a compacted pattern of the tight junction protein claudin-5, and the emergence of specialised perinuclear cell-matrix adhesions. Mechanistically, we find an inverse relationship between claudin-5 levels and the expression of key cell-matrix adhesion proteins zyxin and vinculin *in vitro* and in mice. Finally, we demonstrate that this mechanobiological signature associated with BBB maturation is reversed upon BBB dysfunction after seizures in mice and in human patients with temporal lobe epilepsy. Collectively, our findings establish a novel interplay between mechanoresponsive elements in brain ECs, with implications for BBB stabilisation therapy in epilepsy.

## Introduction

The blood-brain barrier (BBB), formed by highly specialised brain microvessels, controls the composition of the neuronal microenvironment and is vital for neurological function^1^. Endothelial cells (ECs) lining these vessels are constantly exposed to mechanical forces resulting from blood flow-induced shear stress, hydrostatic pressure and extracellular matrix stiffness^2^, which ECs sense and act on to fine-tune BBB integrity. Recent studies have described key molecular players that shape how brain ECs sense such forces^3,4^, including ion channels PIEZO1^5,6^ and TRPV4^7^ as well as structural elements, such as the glycocalyx^8^ and actively suppressed caveolae^9,10^. Yet, the specific mechanisms by which brain ECs then integrate and transmit forces to establish and maintain the BBB are poorly understood.

The two main force-bearing systems in brain ECs are cell-cell junctions (connecting ECs to each other) and cell-matrix adhesions (connecting ECs to their environment), which are both anchored by the actin cytoskeleton^4^. Unlike in peripheral ECs, cell-cell contacts at the BBB feature elaborate tight junctions that limit solute exchange between the blood and brain parenchyma. While the molecular composition of brain endothelial tight junctions is well-characterised^11^, with claudin-5 being the dominant tight junction protein^12,13^, many questions still remain about the precise architecture, mechanics and dynamic regulation of cell-cell contacts at the BBB. Considerably less is known about the arrangement and dynamics of cell-matrix adhesions at the brain vasculature, and how the two mechanoresponsive systems interact with each other to enable barrier formation.

Importantly, the BBB becomes dysfunctional in a wide range of neurological and neuropsychiatric conditions, which is a key driver of pathology in these diseases. Mechanical alterations that lead to BBB leakage and extracellular matrix dysregulation are particularly common in epilepsy^14–18^, a disease defined by recurrent unprovoked seizures that affects more than 50 million people worldwide. We have previously shown that levels of the tight junction protein claudin-5 are negatively associated with epilepsy, with claudin-5 knockdown leading to spontaneous recurrent seizures, severe neuroinflammation and mortality^19^. It is less clear, however, how endothelial cell-matrix adhesion is involved in BBB breakdown, and which mechanical elements in brain ECs might serve as therapeutic targets in disease. As BBB stabilisation has emerged as a promising way to prevent seizure activity^19,20^ and to treat a range of other neurological disorders^21^, there is now a pressing need to understand the dynamic nature of brain EC mechanics, and how they control BBB integrity in both health and disease.

Here, we explore key mechanobiological aspects of BBB maturation and dysfunction. For a robust induction of BBB maturation, we took advantage of our recently developed cARLA method that synergistically enhances barrier tightness and induces complex BBB properties in cultured human ECs^22^. Using the cARLA method, we now reveal a coordinated redistribution of mechanoresponsive elements during human BBB maturation, with a compacted pattern of tight junctions and the emergence of highly specialised perinuclear cell-matrix adhesions. We also describe the first force signature of BBB maturation, and reveal an inverse relationship between tight junction protein levels and the expression of key cell-matrix adhesion proteins *in vitro* and in mice. Finally, we demonstrate a reversal of these mechanobiological properties upon epileptic BBB dysfunction in mice and in brain sections from epilepsy patients. Together, these findings establish a reciprocal regulation of brain vascular integrity by endothelial tight junctions and cell-matrix adhesions in health and epilepsy.

## Results

### Tight junctions and cell-matrix adhesions are markedly redistributed during BBB maturation

Using super-resolution imaging, we first examined how the nano-scale structure of tight junctions changes in human stem cell-derived ECs upon BBB maturation. To induce BBB maturation, we used our well-established cARLA method^22^. As previously demonstrated, a 48-hour treatment with this small molecule cocktail synergistically induces BBB properties in ECs by acting on three signaling pathways that converge on the tight junction protein claudin-5 (**Fig. 1a**).

**Fig. 1.**
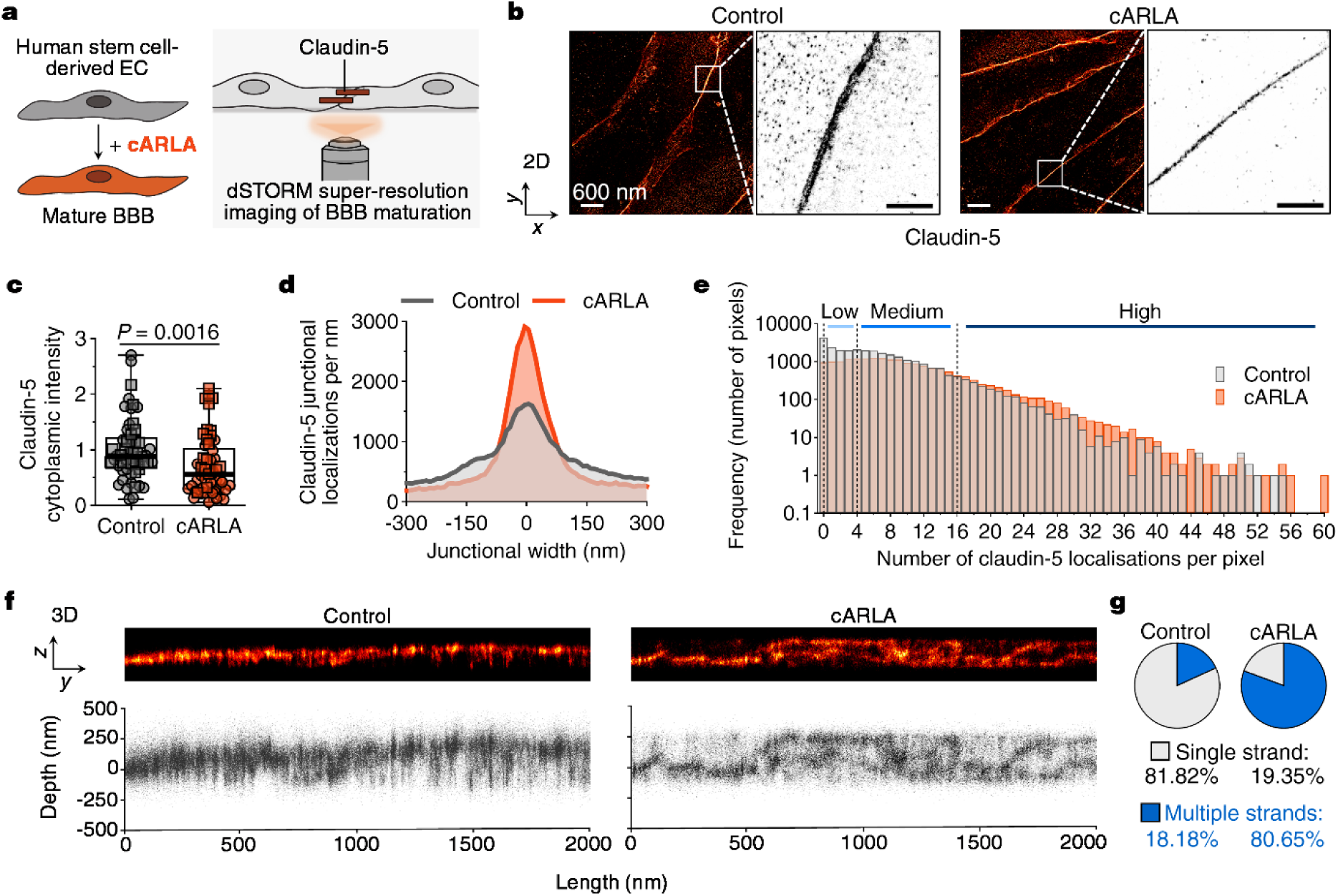
Super-resolution imaging reveals tight junction redistribution during BBB maturation. **a)** Schematic of the experimental setup. The small molecule cocktail cARLA is used to induce BBB maturation. **b)** Representative super-resolution images of the tight junction protein claudin-5 in human stem cell-derived ECs in 2D, as seen from above. Insets show claudin-5 localisations at cell-cell junctions. **c)** Claudin-5 intensity in the cytoplasm. Box: median ± quartiles, whiskers: range. Mann-Whitney test, two-tailed, n=53 cells from two experiments. **d)** Representative density plot showing the number of junctional claudin-5 localisations and tight junction width in ECs during BBB maturation. **e)** Histogram showing the distribution of junctional claudin-5 localisations in ECs, n=43 full-length junctions from two experiments. Three subgroups were created based on localisation density: low (0-4), medium (5-16) and high (17-60 localisations per pixel). **f)** 3D architecture of tight junctions in ECs along the *z*-axis. Representative super-resolution images (*y-z* projection, as seen from the side; top panel) and scatter plots of single claudin-5 localisations are shown (lower panel). **g)** Pie chart showing the proportion of tight junctions composed of single- and multiple strands, n=43 full-length junctions from two experiments.

Three-dimensional direct stochastic optical reconstruction microscopy (3D dSTORM) allowed us to visualize the spatial distribution of single claudin-5 molecules in ECs with nanoscale resolution (**Fig. 1b**). In mature brain ECs, we observed a shift in claudin-5 distribution (**Fig. 1b**), with reduced cytoplasmic (**Fig. 1c**) and increased junctional staining (**Fig. 1d; Supplementary Fig. 1a**) compared to the control group. In addition, the width of claudin-5 staining at cell-cell junctions was markedly reduced upon cARLA treatment (**Fig. 1b,d; Supplementary Fig. 1b**), indicating that mature tight junctions had higher amounts of claudin-5 packed into a more compact space. Specifically, treatment with cARLA increased the number of pixels with high amounts of claudin-5 localisations, and decreased the number of pixels with zero or low amounts of claudin-5 localisations along the length of tight junctions (**Fig. 1e**). In 2D, we also noted that tight junctions at the mature BBB had a less jagged, zig-zaggy morphology (**Supplementary Fig. 1c-d**).

To capture the architecture of tight junctions along the *z*-axis, we performed 3D super-resolution imaging and generated *y/z* (side view) projections of claudin-5 localisations (**Fig. 1f**). Strikingly, while 82% of tight junction strands in the control group were composed of a single strand, this figure was only 19% in the cARLA-treated group (**Fig. 1f,g, Supplementary Fig. 1e,f**). In other words, 81% of tight junctions in mature, cARLA-treated cells were composed of two or more strands of claudin-5 localisations in 3D (**Fig. 1f,g**). Tight junctions strands spanned multiple focal planes and created an intricate meshwork-like structure with ‘kissing points’ between strands (**Fig. 1f**). The 3D structure of strands along the length of junctions was also considerably more complex at the mature BBB (**Supplementary Fig. 1g**).

To better understand changes in claudin-5 spatial distribution, we imaged the underlying cytoskeleton that provides mechanical support for tight junctions (**Fig. 2a**). Filamentous actin was mostly present in stress fibers in ECs under control conditions but was markedly redistributed upon cARLA treatment (**Fig. 2a,b**). The redistribution involved two hotspots: actin was transferred to the cell periphery (cortical actin) and to the area around nuclei (perinuclear actin, **Fig. 2a,b**). Importantly, this suggests that mature brain ECs not only change the way they attach to each other during BBB maturation, but also how they adhere to the extracellular matrix. We found that perinuclear actin was anchored to the extracellular matrix *via* specialised structures, which we termed as perinuclear adhesions (**Fig. 2c**). Perinuclear adhesions stained positive for focal adhesion proteins, such as zyxin, vinculin and the focal adhesion kinase (**Fig. 2c; Supplementary Fig. 2a-f**). Using 3D rendering, we also noted that actin filaments anchored by focal adhesion proteins run over and above nuclei at the mature BBB (**Fig. 2d-g; Supplementary Fig. 2d,e**), possibly to provide extra mechanical stability at the central part of the cell. While the expression of zyxin and vinculin were downregulated by cARLA at the mRNA level (**Fig. 2h**), we did not observe changes in their protein levels as quantified by microscopy (**Fig. 2i**). Rather, perinuclear adhesions specific to the mature BBB were a result of changes in the localisation of focal adhesion proteins, from a uniform distribution to specifically positioned clusters (**Supplementary Fig. 3a-d**). Indeed, while 88% of ECs at the mature BBB had perinuclear adhesions, this figure was only 3% in the control group (**Fig. 2j; Supplementary Fig. 3a-d**). Together, these findings reveal a coordinated redistribution of the endothelial actin cytoskeleton as well as tight junctions and cell-matrix adhesions, key mechanoresponsive structures, during BBB maturation.

**Fig. 2.**
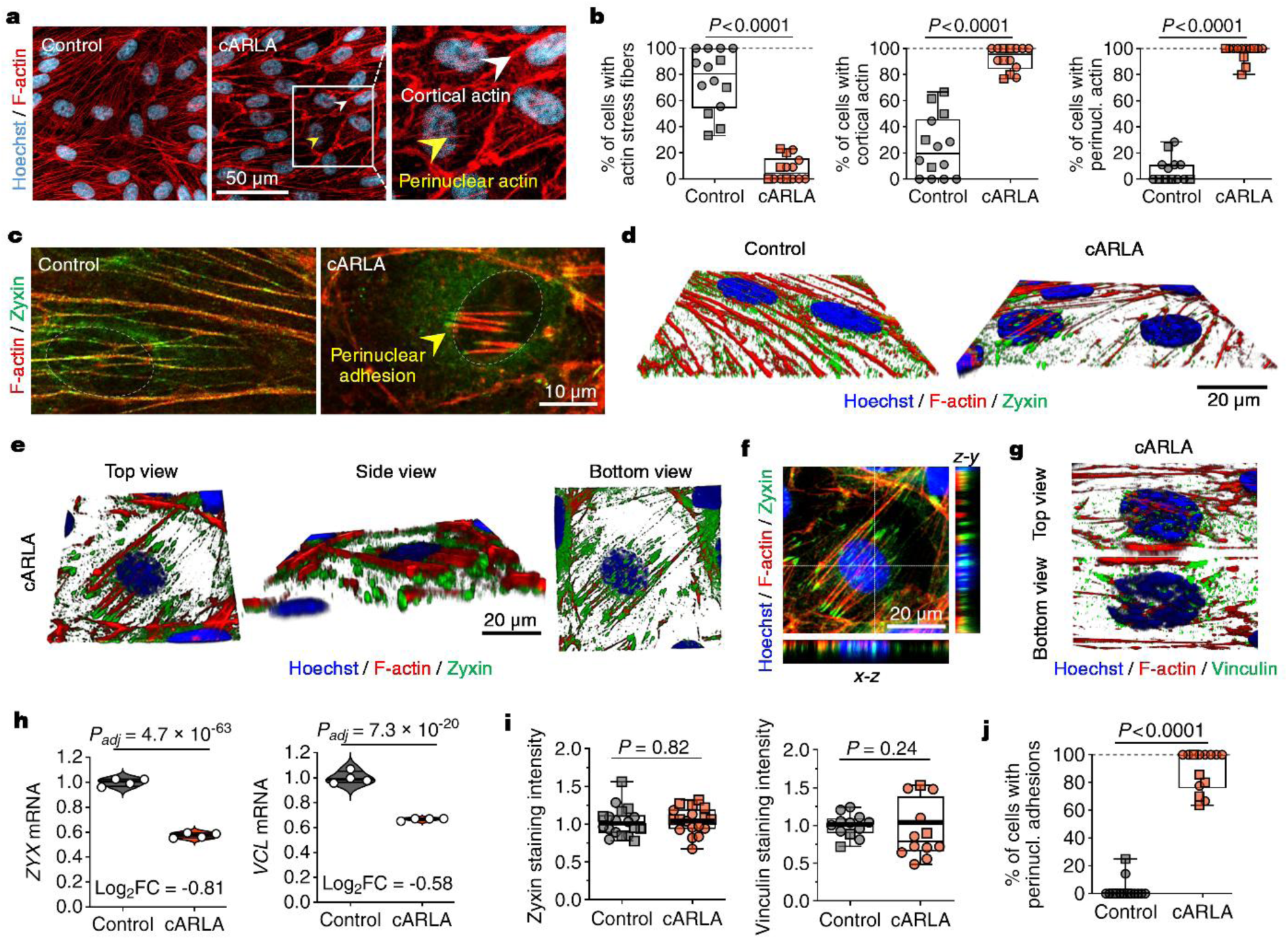
Specialised perinuclear focal adhesions control cell-matrix attachment at the mature BBB. **a)** Representative confocal microscopy images of F-actin in human stem cell-derived ECs during BBB maturation. Insets show the two hotspots of actin rearrangement: cortical-(white arrowhead) and perinuclear actin (yellow arrowhead). **b)** Quantification of the number of cells having actin stress fibers (left panel), cortical actin (middle panel) and perinuclear actin (right panel). Box: median ± quartiles, whiskers: range. Mann-Whitney test, two-tailed, n=14 images from two experiments. **c)** Representative images of F-actin co-stained with the focal adhesion protein zyxin in human stem cell-derived ECs. Yellow arrowheads highlight the emergence of perinuclear adhesions at the mature BBB. **d)** Representative 3D reconstructions of images show the unifom (control) and perinuclear (cARLA-treated) distribution of zyxin in ECs. **e)** 3D reconstructions of images in cARLA-treated mature ECs from multiple angles, and **f)** *x-z* and *y-z* projections in 2D highlight the spatial organisation of F-actin and zyxin in perinuclear adhesions. **g)** Representative 3D reconstructions from cARLA-treated mature ECs showing a similar organisation of the focal adhesion protein vinculin in perinuclear adhesions. **h)** Changes in *ZYX*/zyxin and *VCL*/vinculin mRNA levels during BBB maturation as determined by RNA-sequencing. Log_2_FC: log_2_ (fold change). *P*-values were adjusted for multiple comparisons (FDR; Benjamini-Hochberg method), n=4 cell culture inserts per group. **i)** Changes in zyxin and vinculin staining intensity as determined by confocal microscopy. Box: median ± quartiles, whiskers: range. Unpaired t-test, two-tailed, n=14 images (zyxin) and n=12 images (vinculin) per group from two experiments. **j)** Quantification of the number of of cells having perinuclear adhesions. Box: median ± quartiles, whiskers: range. Mann-Whitney test, two-tailed, n=14 images from two experiments.

### A force signature of BBB maturation over time

After quantifying morphological changes in force-bearing endothelial structures, we were curious how such changes create a functional output during BBB maturation. To explore this, we measured the force signature of BBB maturation using fluidic force microscopy (FluidFM). In FluidFM, a specialised hollow cantilever probe (**Fig. 3a**) is used to engage and detach individual ECs from a monolayer using negative pressure and force-controlled cantilever liftup (**Fig. 3b,c**). Strikingly, when applying such negative pressure that successfully detached an individual EC from control monolayers in 95% of cases (19/20 trials), we could only detach ECs from cARLA-treated monolayers with a 12.5% success rate (5/40 trials, **Fig. 3d**). This indicates that individual ECs are more tightly embedded in monolayers at the mature BBB. We wondered if this was the result of increased cell-cell adhesion, cell-matrix adhesion or both.

**Fig. 3.**
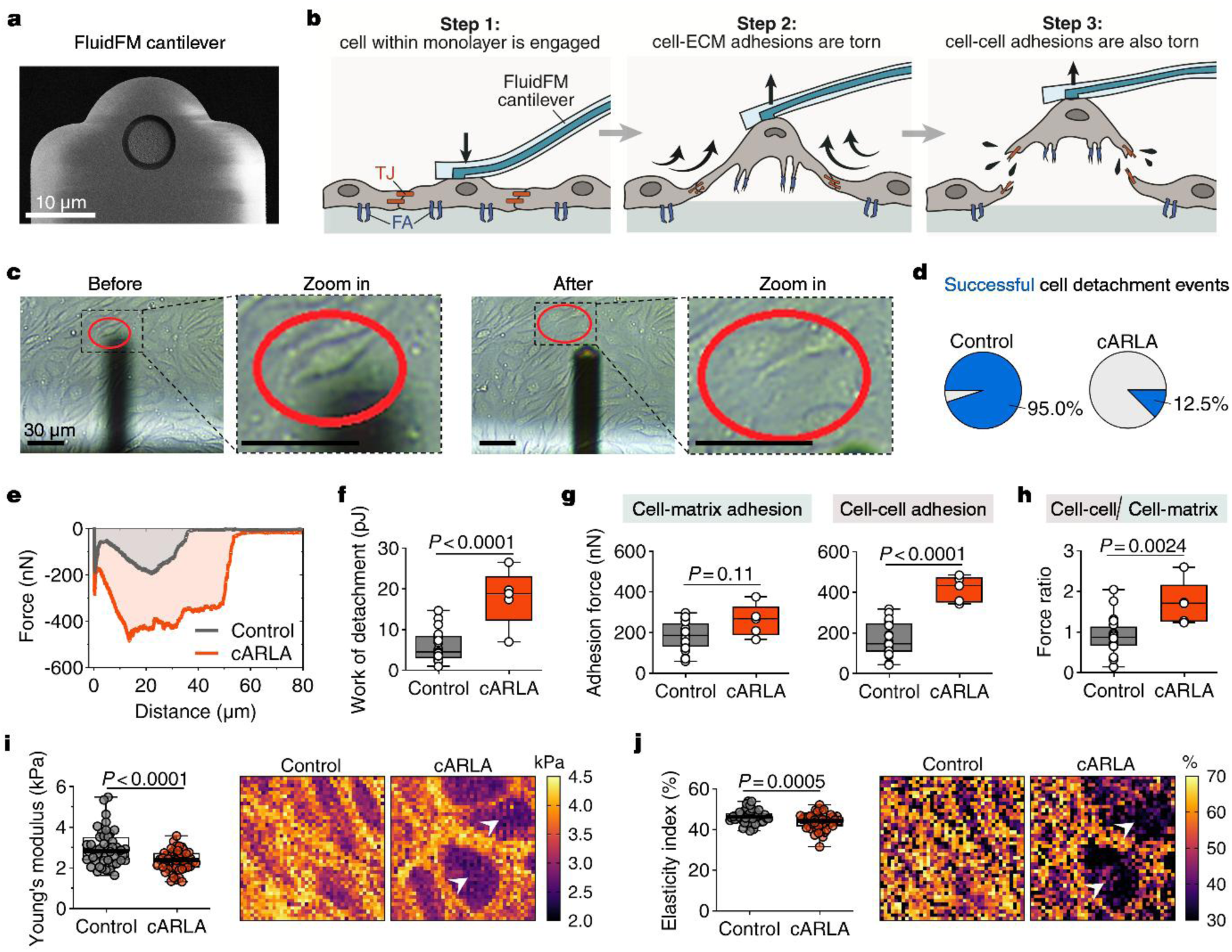
A force signature of BBB maturation. **a)** Image of the specialised hollow cantilever with circular opening (microprobe) used in fluid force microscopy (FluidFM). **b)** Schematic of the FluidFM measurement process. After a cell is engaged with the cantilever using negative pressure, a force-controlled lift-up protocol is applied to detach that cell from the monolayer. First, cell-matrix adhesions are torn. Cell-cell adhesions are torn in a subsequent step, and the cell is then detached. TJ: tight junction, FA: focal adhesion. **c)** The process of cell detachment by FluidFM is shown using real-time phase-contrast microscopy in human stem cell-derived EC monolayers. The red circle highlights the cell that was engaged and detached. **d)** Pie chart showing the proportion of successful cell detachment events using the same negative pressure; n=21 (control) and n=40 (cARLA) total cell detachment events. **e)** Representative force-distance curves of the cell detachment process in control and mature ECs. The first minimum corresponds to cell-matrix andhesions, the second minimum corresponds to cell-cell adhesions. **f)** Differences in total work of detachment (area under the curve) between control and cARLA-treated ECs based on successful cell detachment events. **g)** Adhesion forces corresponding to cell-matrix (left panel) and cell-cell adhesion (right panel) upon BBB maturation. **h)** Ratio of cell-cell vs. cell-matrix adhesion from the previous panel. In all box plots, box: median ± quartiles, whiskers: range, unpaired t-test, two-tailed. In panels **f,g** and **h**: n=20 cells (control) and n=5 cells (cARLA) per group from two experiments. In panels **i** and **j**: n=47 images per group from four experiments.

Focusing on successful cell detachment events, we generated force-distance curves of the detachment process (**Fig. 3e; Supplementary Fig. 4**). These curves had a characteristic shape with two local minima (dips), the first dominated by cell-matrix adhesion forces and the second one dominated by cell-cell adhesion forces^23^ (**Fig. 3b,e**). In line with the observed differences in cell detachment efficiency, the overall work of detachment (area under the curve) was significantly higher at the mature BBB (**Fig. 3f**). Importantly, this was a result of increased cell-cell, but not cell-matrix, adhesion (**Fig. 3g**). While control cells exhibited approximately equal forces at cell-cell and cell-matrix adhesions, this balance was shifted towards cell-cell adhesions in cARLA-treated ECs (**Fig. 3h**) without a reduction in forces on cell-matrix adhesions (**Fig. 3g**). We also validated these findings using atomic force microscopy (AFM; **Fig. 3i,h**; **Supplementary Fig. 5a-d**). Compared to control ECs having a more uniform force distribution profile, mature brain ECs were softer and more plastic at the area above nuclei but harder and more elastic at cell-cell junctions (**Fig. 3i,h**), confirming the redistribution of forces. Of note, cell height remained unchanged upon cARLA treatment (**Supplementary Fig. 5d**).

Next, we investigated how this force redistribution is orchestrated over time. We used resonant waveguide grating (RWG), a label-free and high-throughput biophysical technique, to measure the real-time kinetics of cell-matrix adhesion during BBB maturation (**Fig. 4a; Supplementary Fig. 6a**). RWG quantifies the resonant wavelength shift of light caused by cellular structures at the basal 150 nm part of an EC monolayer, which is directly proportional to the surface area covered by cell-matrix adhesions and their local density^24^ (**Fig. 4a**). During the early phase of BBB maturation, cARLA-treated ECs induced a positive wavelength shift that returned to baseline within 2 hours (**Fig. 4b**). By contrast, at mid (24 h) and late (48 h) timepoints after cARLA treatment, we measured progressively more negative wavelength shifts compared to both baseline and the control group (**Fig. 4b; Supplementary Fig. 6a,b**). This indicates a gradual decrease in the surface area covered by cell-matrix adhesions during BBB maturation. Indeed, while the early phase of BBB maturation (0-2 h) was dominated by an evenly distributed pattern of focal adhesions, a complete remodeling of cell-matrix contacts was apparent at mid- and late phases (**Fig. 4c; Supplementary Fig. 6c-f**). At 24 and 48 hours after cARLA treatment, 46% and 88% of ECs had zyxin-containing perinuclear adhesions, respectively (**Fig. 4d**). A similar tendency was seen for perinuclear actin filaments (**Fig. 4e**). Importantly, perinuclear actin and perinuclear adhesions were almost entirely absent at early timepoints and in the control group at 48 hours (**Fig. 4c-e; Supplementary Fig. 6d-f**), further confirming that these structures are specific to the mature BBB. Collectively, we propose a mechanobiological signature of BBB maturation in which progressively more force is placed on cell-cell adhesions, while a constant amount of force is placed on progressively fewer, highly specialised perinuclear cell-matrix adhesions over time.

**Fig. 4.**
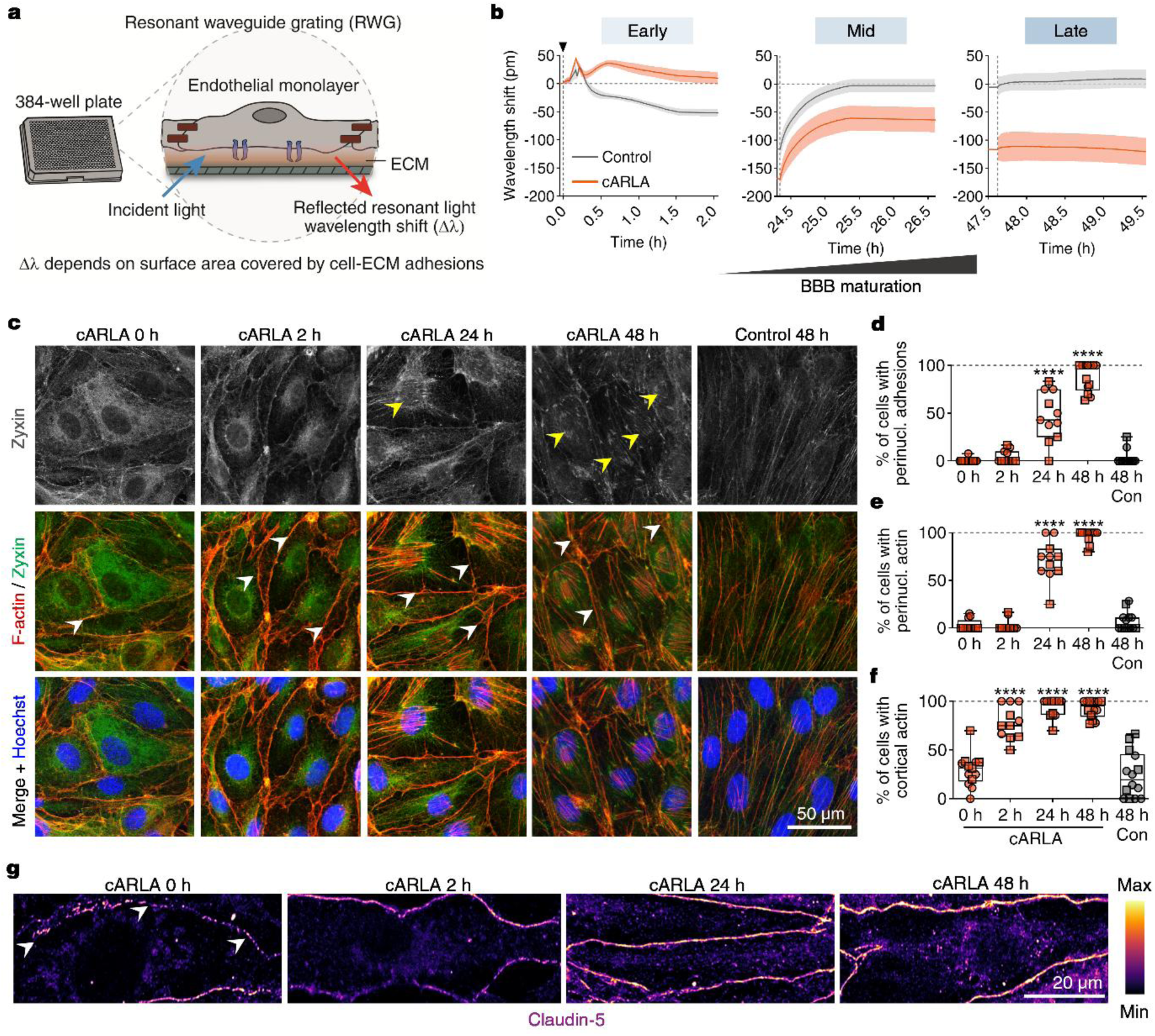
Kinetics of force redistribution during BBB maturation. **a)** Schematic of experimental setup. Resonant waveguide grating measures the resonant wavelength shift (Δλ) caused by cellular structures at the basal part of cells, which is directly proportional to the surface area covered by cell-matrix adhesions. ECM: extracellular matrix. **b)** Wavelength shift kinetics of human stem cell-derived brain ECs at early, mid and late timepoints of *in vitro* BBB maturation. **c)** Representative confocal microscopy images of zyxin and F-actin distribution upon cARLA treatment over time. Yellow arrowheads in the top row point to zyxin-containing perinuclear adhesions. White arrowheads in the middle row point to cortical actin structures. **d)** Quantification of the number of cells having perinuclear adhesions, **e)** perinuclear actin and **f)** cortical actin during BBB maturation. Note that the emergence of cortical actin precedes perinuclear actin and perinuclear adhesions. In all three panels, box: median ± quartiles, whiskers: range. One-way ANOVA compared to the 0h cARLA group, \*\*\*\**P*<0.0001, n=11-14 images from two experiments. **g)** Claudin-5 continuity at cell-cell junctions upon cARLA treatment over time. White arrowheads indicate discontinous junctions. Note the rapid increase in claudin-5 staining intensity and continuity at cell-cell junctions.

### BBB tightness negatively regulates endothelial cell-matrix adhesion *via* claudin-5

While perinuclear actin- and adhesions emerged during mid-to-late phases of *in vitro* BBB maturation (**Fig. 4c-e**), cortical actin was already strongly present at the cell periphery 2 hours after cARLA treatment (**Fig. 4c,f**). This timing coincided with rapid changes in claudin-5 distribution and continuity at tight junctions (**Fig. 4g; Supplementary Fig. 7a**), suggesting that changes to cell-cell junctions precede the rearrangement of cell-matrix adhesions during BBB maturation. Additionally, trypsin-mediated cleavage of cell-cell junctions resulted in an immediate positive wavelength shift in cARLA-treated ECs, indicating a rapid increase in the surface area covered by cell-matrix contacts (**Supplementary Fig. 7b-d**). Based on these observations, we hypothesised that junctional tightness can directly influence the state of cell-matrix adhesions at the BBB.

To test this, we generated brain EC populations with different levels of junctional tightness by stably overexpressing (OE) or knocking out (KO) claudin-5 from mouse bEnd.3 cells (**Fig. 5a,b**). Using RNA-sequencing, we indeed identified a core set of cell-matrix adhesion genes that inversely followed claudin-5 levels across OE, KO and mock-transfected brain EC populations (**Fig. 5c,d**). These genes encode focal adhesion proteins (*Zyx/*zyxin*, Vcl/*vinculin*, Vasp*), key mediators of integrin signalling (*Itgav, Itgb1, Epha2*), extracellular matrix components (*Col4a1, Col4a2, Hspg2*) and stress fiber-associated proteins (*Micall2, Pdlim1, Synpo*; **Fig. 5c,d**). Notably, all of these genes are relatively depleted from brain ECs and are more enriched in peripheral ECs in mice (**Supplementary Fig. 8**).

**Fig. 5.**
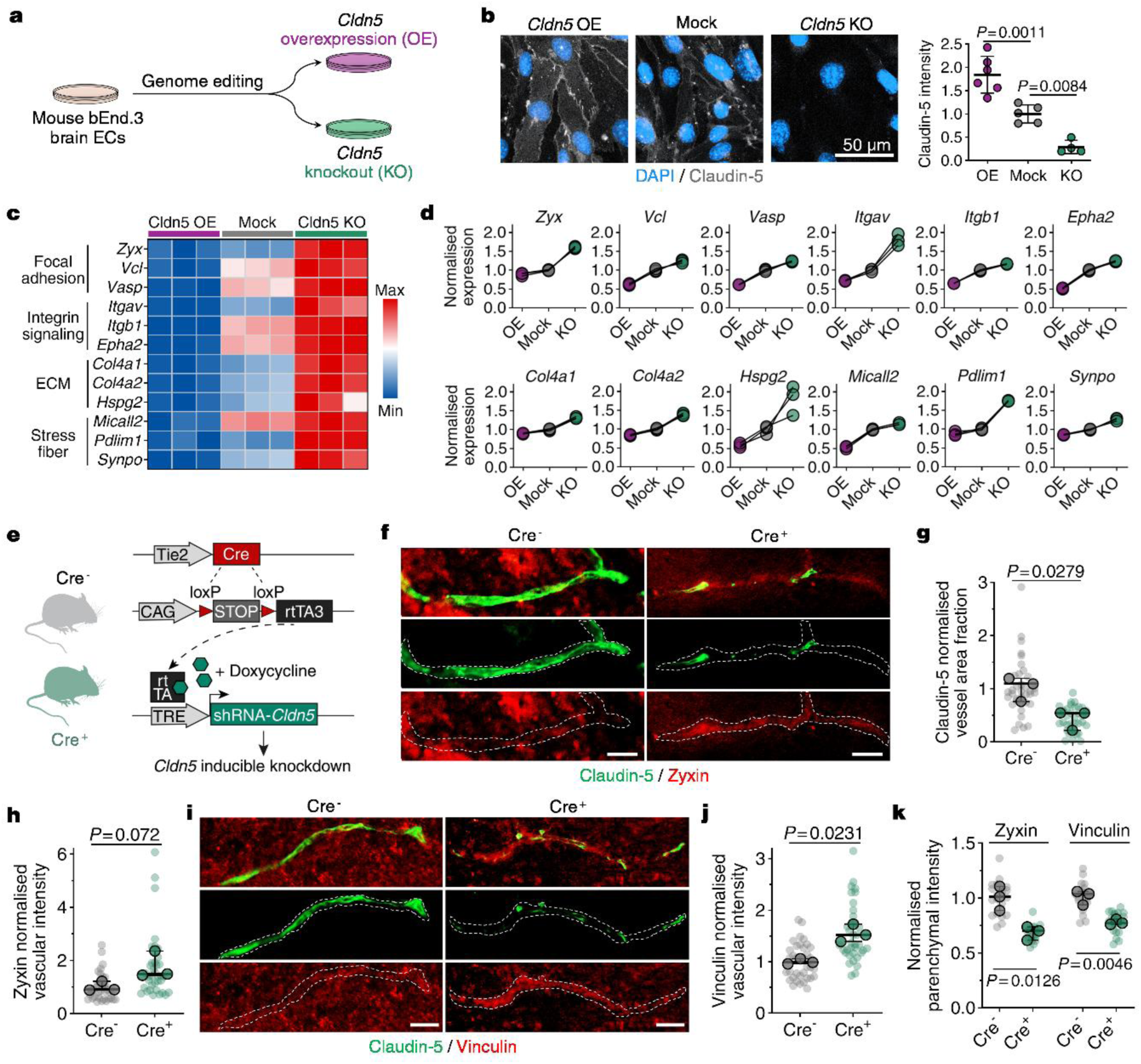
BBB tightness negatively regulates endothelial cell-matrix adhesion *via* claudin-5. **a)** Schematic of claudin-5 overexpression (OE) and knockout (KO) strategy *in vitro*. **b)** Claudin-5 immunostaining in *Cldn5* OE, KO and mock-transfected cell populations (left panel). Quantification (right panel): mean ± SD, one-way ANOVA, n=4-7 images per group. **c)** Min-max scaled heat map and **d)** Slope chart showing a core set of cell-matrix adhesion genes that inversely follow claudin-5 levels *in vitro* as determined by RNA-sequencing. For each gene, between-group FDR-corrected *P*-values (Benjamini-Hochberg method) < 0.05; n=3 cell culture inserts per group. ECM: extracellular matrix. **e)** Schematic of the *Cldn5* knockdown mouse model used. Upon doxycycline supplementation in adult mice, an inducible shRNA against *Cldn5* suppresses claudin-5 levels in littermates positive for an endothelial cell-specific Cre recombinase (Cre^+^). Cre^-^ littermates were used as controls. **f)** Immunohistochemistry of claudin-5 and zyxin in mouse brain sections with or without *Cldn5* knockdown. Dashed lines outline blood vessels. Scale bar: 15 µm. **g)** Quantification of claudin-5 and **h)** vessel-associated zyxin levels. For both panels: unpaired t-test with Welch’s correction, two-tailed, n=36 blood vessels from 3 mice per group. **i)** Immunohistochemistry of claudin-5 and vinculin in mouse brain sections with or without *Cldn5* knockdown. Dashed lines outline blood vessels. Scale bar: 15 µm. **j)** Quantification of vessel-associated vinculin levels. unpaired t-test with Welch’s correction, two-tailed, n=36 blood vessels from 3 mice per group. **k)** Quantification of parenchymal zyxin and vinculin levels. Unpaired t-test with Welch’s correction, two-tailed, n=20 equally sized parenchymal areas from 3 mice per group. In panels **g**-**h** and **j-k**, lighter dots indicate individual blood vessels and darker dots indicate means from each animal. Statistical tests were performed on means from each animal.

Using an inducible *Cldn5* knockdown mouse model^25^ (**Fig. 5e**), we validated findings at the protein level for zyxin and vinculin. In these adult mice, a doxycycline-inducible shRNA against *Cldn5* allows the suppression of claudin-5 levels over time in littermates positive for an EC-specific Cre recombinase (**Fig. 5e**). After doxycycline supplementation, Cre^+^ mice had 57% lower claudin-5 protein levels than Cre^-^ littermate controls (**Fig. 5f,g**). In line with our *in vitro* data on claudin-5 KO bEnd.3 cells, we observed a 1.76-fold increase in vessel-associated zyxin, with a strong trend for statistical significance (**Fig. 5f,h**) and a 1.55-fold increase in vessel-associated vinculin (**Fig. 5i,j**) in Cre^+^ animals compared to Cre^-^ controls. Notably, Cre^-^ control mice showed mainly parenchymal, but not vascular, zyxin and vinculin staining (**Fig. 5f,I**), both of which were reduced in Cre^+^ littermates (**Fig. 5k**). This indicates increased vascular and decreased parenchymal levels of zyxin and vinculin upon *Cldn5* knockdown. Taken together, our results show that endothelial cell-matrix adhesions inversely follow junctional tightness, and specifically claudin-5 levels, at the BBB *in vitro* and in mice. Given the key role of claudin-5 in regulating BBB integrity, we wondered how endothelial cell-matrix adhesion is orchestrated during pathological BBB disruption.

### Increased endothelial cell-matrix adhesion in epileptic BBB dysfunction in mice and humans

We have previously shown that reduced tight junction integrity and reduced claudin-5 levels are strongly associated with epilepsy in mice and in human patients^19^. To assess if brain endothelial cell-matrix adhesions are altered in epilepsy, we reanalysed a publicly available RNA-sequencing dataset on brain ECs from a kainic acid-induced epilepsy mouse model^26^ (**Fig. 6a**). Indeed, the top pathways in brain ECs acutely induced by seizures were related to cell-substrate adhesion, focal adhesions and actin stress fiber formation (**Fig. 6a; Supplementary Table 1**). Importantly, 11 out of 12 genes from our cell-matrix adhesion gene module (**Fig. 5c,d**) were highly upregulated in seizure-bearing mice (**Fig. 6b**). The regulation of 8 of these genes (*Zyx, Vcl, Vasp, Itgav, Epha2, Col4a1, Col4a2, Synpo*) was also highly consistent between 3 different RNA-sequencing datasets from different biological contexts (**Supplementary Fig. 9a-c**). Our top hits, zyxin/*Zyx* and vinculin/*Vcl* inversely followed *Cldn5* levels (**Fig. 5d**), were downregulated by cARLA during BBB maturation (**Fig. 2h**; **Supplementary Fig. 9a**), and were upregulated in epilepsy in mice by 2.6 and 5-fold, respectively (**Fig. 6b**).

**Fig. 6.**
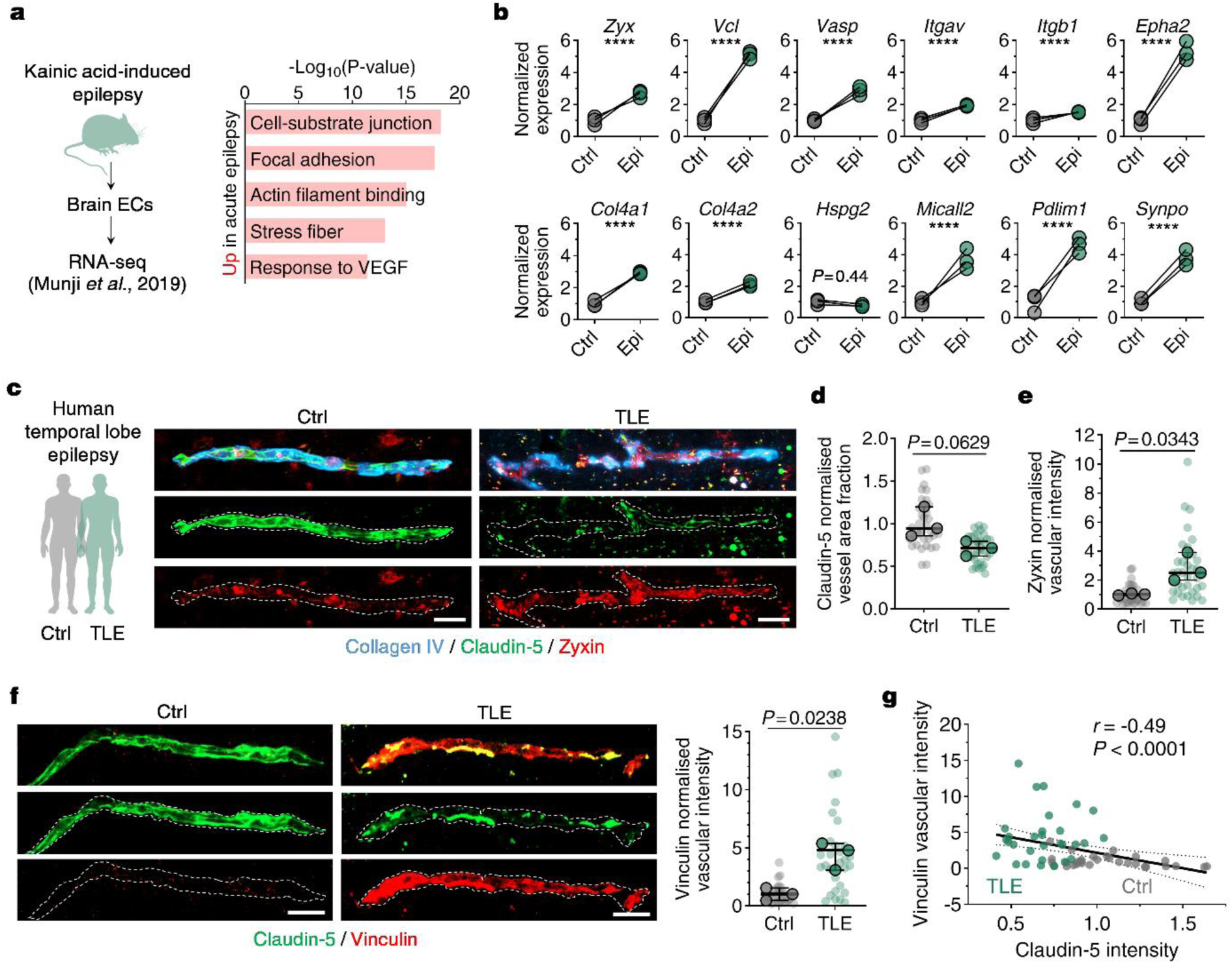
Increased endothelial cell-matrix adhesion in epileptic BBB dysfunction in mice and humans. **a)** RNA-sequencing analysis of mouse brain ECs in epilepsy. Left panel: schematic of the experimental setup. Right panel: top upregulated pathways in brain ECs in acute epilepsy as reanalysed from Munji *et al*.^26^ **b)** Analysis of our core set of 12 cell-matrix adhesion genes that inversely follow claudin-5 levels in the acute epilepsy dataset. Differential expression analysis by DESeq2, FDR-corrected *P*-value (Benjamini-Hochberg method), *\*\*\*\*P_adj_*<0.0001, n=3 mice per group. **c)** Immunohistochemistry of collagen type IV, claudin-5 and zyxin in human brain sections from autopsy controls or patients with temporal lobe epilepsy (TLE). Dashed lines outline blood vessels. Scale bar: 20 µm. **d)** Quantification of claudin-5 and **e)** vessel-associated zyxin levels. Unpaired t-test with Welch’s correction, two-tailed, n=36 blood vessels from 3 patients per group. **f)** Immunohistochemistry of claudin-5 and vinculin in human brain sections from autopsy controls or patients with temporal lobe epilepsy (TLE). Dashed lines outline blood vessels. Scale bar: 20 µm. Quantification (right panel): unpaired t-test with Welch’s correction, two-tailed, n=30 blood vessels from 3 patients per group. In panels **d-f**, lighter dots indicate individual blood vessels and darker dots indicate means from each patient. Statistical tests were performed on means from each patient. **g)** Correlation analysis of vessel-associated vinculin and claudin-5 levels in human brain sections. Spearman correlation, two-tailed, n=60 datapoints from both control (grey) and TLE (green) groups. The line of best fit (continuous line) and 95% confidence intervals (dashed lines) from a simple linear regression are shown.

With a proof-of-concept approach, we assessed zyxin and vinculin protein distribution in resected human brain tissue from patients with temporal lobe epilepsy (TLE, n=3, **Supplementary Table 2**) and matched autopsy controls (Ctrl, n=3; **Supplementary Table 3**). In agreement with previously reported values^19^, claudin-5 protein levels were reduced by ∼30% in TLE patients, with a strong trend for statistical significance (**Fig. 6c,d**). Strikingly, we observed a 2.8-fold increase in vessel-associated zyxin levels in TLE patients (**Fig. 6c,e**). In addition, while zyxin localised to well-positioned clusters within brain microvessels in the control group, it was more uniformly distributed in vessels from TLE patients **(Fig. 6c**), inversely mirroring our findings on *in vitro* BBB maturation (**Fig. 2c,j**). Similarly, vinculin was not associated with blood vessels in the control group but its vascular localisation and protein levels were elevated in the TLE group by 4.4-fold (**Fig. 6f**). Added to this, vessel-associated vinculin levels inversely correlated with claudin-5 levels in both human (*r*=-0.49, *P*<0.0001; **Fig. 6g**) and mouse brain sections (*r*=-0.57, *P*<0.0001; **Supplementary Fig. 10a**). Interestingly, although zyxin protein levels were highly increased after BBB disruption, we found no correlation between vascular zyxin and claudin-5 levels in mouse and human brain sections (**Supplementary Fig. 10b,c**). Collectively, these results indicate that endothelial cell-matrix adhesion is markedly increased in epileptic BBB dysfunction in mice and in human patients. Our data also suggest that vessel-associated zyxin and vinculin respond differently to changes in BBB tightness and claudin-5 levels.

## Discussion

In this study, we revealed i) how brain endothelial force-bearing structures and mechanical forces are redistributed during BBB maturation, ii) described how tight junctions and cell-matrix adhesions reciprocally control barrier integrity at the molecular level and iii) demonstrated the pathological relevance of these findings in epileptic BBB dysfunction.

First, we used super-resolution microscopy to image nanoscopic changes in tight junction complexes during BBB maturation. Two recent studies have pioneered this field^27,28^, however, questions still remain about the 3D architecture of tight junctions at the BBB. The strength of our approach compared to previous studies is the use of 3D dSTORM imaging with nanoscale axial resolution. This allowed us to discover that higher amounts of claudin-5 protein are packed into a tighter space in 2D, and that this corresponds to an intricate pattern of multiple intertwining tight junction strands in 3D. Our images show, for the first time using light microscopy, a 3D pattern of tight junctions similar to those in freeze-fracture electron microscopy studies in brain ECs^29–32^. These observations of a meshwork-like structure with kissing points between strands suggest an additional layer of complexity in the regulation of tight junctions at the BBB, in accordance with recent seminal work on the sophisticated organisation of epithelial tight junctions^33,34^.

In parallel to changes in tight junction structure, we observed major rearrangements in the underlying actin cytoskeleton and cell-matrix adhesions in ECs during BBB maturation, with hotspots at the cell periphery and around nuclei. Intriguingly, actin around the nucleus has been described as an important regulator of ultrafast ‘outside-in’ mechanotransduction and cell shape in fibroblasts^35–37^, and similar structures have recently been reported in lymphatic ECs subjected to high levels of mechanical stretch^38^. We found that perinuclear actin was anchored to the extracellular matrix *via* zyxin and vinculin-containing structures, which we termed as perinuclear adhesions. Perinuclear adhesions were specific to the mature BBB, and we hypothesize that they are key to i) allow force redistribution in ECs to support increased loads on tight junctions, while ii) providing balance and stability for ECs.

Our study provides the first comprehensive biophysical characterisation of BBB maturation, which we performed in a time-dependent manner, using three different techniques. Our FluidFM and RWG measurements suggest a model in which progressively more force is placed on EC-EC contacts, while a constant amount of force is placed on progressively fewer, highly specialised perinuclear cell-matrix adhesions over time. Importantly, this indicates that by rearranging and concentrating forces to specific subcellular locations, mature brain ECs can achieve the same level of cell-matrix adhesion as control ECs while also creating a tighter paracellular barrier. These functional biophysical measurements are supported by our morphological observations on claudin-5, the actin cytoskeleton and focal adhesions proteins. As a final validation, we used AFM to combine functional and morphological outputs by mapping force measurements to subcellular locations in ECs^39–41^. Indeed, our AFM images show a shift from uniform to polarised force distribution upon BBB maturation, with regions above cell-cell junctions being harder and more elastic, while the region above the nucleus being softer, more plastic and more indentable in mature brain ECs.

How do the major force-bearing systems in brain ECs interact with each other? In peripheral ECs, cell-matrix adhesions can directly influence barrier integrity^42,43^. However, it was unclear whether the same was true for brain ECs having a significantly tighter barrier with highly specialised cell-cell junctions, and whether barrier tightness could influence cell-matrix adhesions. A recent study has provided an example of how one mechanoresponsive system can act on another, and thereby regulate BBB integrity^44^. In their work, Ayloo and colleagues described that pericytes can physically pull on ECs *via* basement membrane vitronectin-integrin contacts, which suppresses caveolae formation and non-specific transcytosis at the BBB^44^. We have now demonstrated that junctional tightness directly controls the state of cell-matrix adhesions at the BBB *via* claudin-5, which we validated using genetically engineered ECs *in vitro* and in an inducible claudin-5 knockdown mouse model^25^. Mechanistically, the timeline that emerges from our functional RWG measurements and morphological observations suggests that tight junctions rapidly control cell-matrix adhesions in a cytoskeleton-linked process, presumably mediated by protein phosphorylation events. Such rapid structural changes are then ‘locked in’ by gene expressional changes occuring at later timepoints to fully establish barrier integrity.

Finally, we have now shown that in parallel to a reduction in BBB integrity, endothelial cell-matrix adhesion is increased in epilepsy in mice and human TLE patients. The next logical steps for future studies will be to i) validate this proof-of-concept finding on a larger sample, and ii) to establish whether an increased brain endothelial cell-matrix adhesion is part of the pathomechanism of epileptic BBB dysfunction ‒ or rather, a compensatory protective mechanism that is beneficial for ECs. Inhibition/loss of the focal adhesion kinase has been shown to enhance barrier function in peripheral blood vessels^45,46^, and to reduce tumour-induced vascular permeability in glioma^47^. These findings, together with our own observations, raise the idea of modulating EC focal adhesions to regulate BBB integrity in epilepsy. As up to one-third of epilepsy patients are refractory to traditional anti-seizure medications^48^, and as brain microvascular stabilisation has recently emerged as a promising way to prevent seizure activity^19,20^, exploring such novel targets is a top priority. Intriguingly, recent omics studies have indicated that the dysregulation of brain EC focal adhesions and extracellular matrix are not unique to epilepsy but are a shared feature of multiple neurological disorders, together with BBB dysfunction and claudin-5 loss^26,49–52^. Therefore, we anticipate that future studies will focus on understanding the dynamics of brain endothelial cell-matrix adhesion as well as explore its modulation as a novel form of microvascular stabilisation therapy against a wide range of neurological and neuropsychiatric conditions.

In conclusion, our study establishes a reciprocal regulation between brain endothelial tight junctions and cell-matrix adhesions during BBB maturation and dysfunction. Together, these findings provide fundamental mechanobiological insight into the regulation of BBB integrity in health and disease, with broad implications for improved benchmarking of *in vitro* BBB models and for brain vascular stabilisation in neurological disorders.

## Materials and methods

### Human brain tissue

Experiments involving human brain tissue were performed in accordance with the Declaration of Helsinki. Studies were approved by the Ethics (Medical Research) Committee of Beaumont Hospital, Dublin, Ireland (ethics code: 20/59), and informed consent was obtained from all participants. Surgically resected brain tissue was obtained from temporal lobe epilepsy (TLE) patients (n=3) at Beaumont Hospital. Post-mortem hippocampal sections of non-diseased control brains (n=3) were obtained from the Stanley Medical Research Institute. Patient demographics can be found in **Supplementary Tables 2-3**.

### Animal experiments

Animal experiments were performed in accordance with the EU Directive 2010/63/EU about animal protection and welfare. Protocols were reviewed and approved by the Research Ethics Committees of Trinity College Dublin and/or the Royal College of Surgeons in Ireland under licenses from the Department of Health (HPRA: AE19136/PO80 and/or AE19127/P057), Dublin, Ireland. The generation of doxycycline-inducible *Cldn5* knockdown mice was described previously^25^. Doxycycline was administered in the drinking water of mice (2 mg/mL in 2% sucrose solution). Mice were kept on doxycycline until they experienced their first seizure, then were sacrificed for tissue collection. Mice were housed under standard conditions (18–23 °C, 12 h light/dark cycle, 40-50% humidity) with regular rodent chow diet and water available *ad libitum*. Mice were bred on-site at Trinity College Dublin onto a C57BL/6J background and used at 8–14 weeks of age for experiments.

### Derivation and culture of human stem cell-derived endothelial cells

CD34^+^ hematopoietic stem cells were isolated from human umbilical cord blood and differentiated towards endothelial cells as previously described^53^. Research was performed according to the principles of the Declaration of Helsinki. Informed consent was obtained from the parents of donors and protocols were approved by the French Ministry of Higher Education and Research (CODECOH DC2011-1321). Endothelial cells (ECs) were derived from CD34^+^ stem cells in endothelial cell growth medium (EGM; Lonza cat# CC-3162) supplemented with 20% fetal calf serum (FCS; Sigma-Aldrich cat# F7524) and 50 ng/mL vascular endothelial growth factor (VEGF165; PeproTech, cat# 100-20). Differentiated ECs were cultured in endothelial cell culture medium (ECM; ScienCell, cat# 1001) supplemented with 5% fetal bovine serum (FBS; Sigma-Aldrich, cat# F4135), 1% endothelial cell growth supplement (ECGS; ScienCell, cat# 1052) and 50 μg/mL gentamicin (Sigma-Aldrich, cat# G1397). ECs were kept in a humidified incubator at 37 °C with 5% CO_2_, and were used between passage numbers 6-7 in all experiments.

### Induction of blood-brain barrier maturation by cARLA

Blood-brain barrier (BBB) properties were induced in human stem cell-derived ECs as previously described^22^. Confluent EC monolayers were treated with the small molecule cocktail cARLA, which is composed of 8-(4-chlorophenylthio)adenosine 3′,5′-cyclic monophosphate sodium salt (cPT-cAMP; Sigma-Aldrich, cat# C3912, 250 μM), Ro 20-1724 (Sigma-Aldrich, cat# 557502, 17.5 μM), lithium chloride (LiCl; Sigma-Aldrich, cat# L9650, 3 mM) and A83-01 (Tocris, cat# 2939, 3 μM). ECs were treated with cARLA for 48 h from the luminal (‘blood side’) compartment.

### Immunocytochemistry

Human stem-cell derived ECs were cultured in glass-bottom chamber slides (Nunc Lab-Tek II, 4-well, Thermo Fisher cat# 154526) or glass cover slips coated with a mixture of collagen IV (Sigma-Aldrich, cat# C5533, 100 µg/mL) and fibronectin (Sigma-Aldrich, cat# F1141, 25 µg/mL) at a seeding density of 6.0×10^4^ cells/cm^2^. Confluent monolayers were treated with cARLA or control medium for 48 h to induce BBB maturation. After 48 h treatment, the medium was removed and ECs were fixed with a 1:1 mixture of ice cold methanol-acetone solution for 2 min (claudin-5 staining) or with 3% paraformaldehyde (focal adhesion proteins) for 15 min at room temperature. For focal adhesion staining, a permeabilization step with 0.2% Triton X-100 in PBS for 10 min at room temperature preceded blocking. Cells were blocked with 3% BSA in PBS for 1 h at room temperature and were subsequently incubated with primary antibodies diluted in blocking buffer overnight at 4 °C. Cells were then incubated with secondary antibodies and the nuclear counterstain Hoechst 33342 (Thermo Fisher, cat# H1399, 1 μg/mL) diluted in PBS for 1 h at room temperature, protected from light. Antibodies/labeling probes used are listed in **Supplementary Table 4**. Between each step, cells were washed three times in PBS. Chamber slides were coverslipped using Fluoromount-G mounting buffer (Southern Biotech, cat# 0100-01) and imaged as described below.

### dSTORM super-resolution microscopy

Human stem-cell derived ECs were cultured on glass cover slips coated with collagen IV and fibronectin, treated with cARLA or control medium for 48 hours, fixed in ice cold methanol-acetone 1:1 solution and stained for claudin-5 as described above. Antibodies used are listed in **Supplementary Table 4**. Samples were mounted onto a microscope slide, then 3D dSTORM experiments were conducted in GLOX switching buffer^54^. The imaging buffer (pH 7.4) was an aqueous solution diluted in PBS containing the GluOx enzymatic oxygen scavenging system, 2000 U/mL glucose oxidase (Sigma-Aldrich, cat# G2133-50KU), 40, 000 U/mL catalase (Sigma-Aldrich, cat# C100), 25 mM potassium chloride (Sigma-Aldrich, cat# 204439), 22 mM tris(hydroxymethyl)aminomethane (Sigma-Aldrich, cat# T5941), 4 mM tris(2-carboxyethyl)phosphine (TCEP) (Sigma-Aldrich, cat# C4706) with 4% (w/v) glucose (Sigma-Aldrich, cat# 49139), and 100 mM β-mercaptoethylamine (MEA, Sigma-Aldrich, cat# M6500). 3D dSTORM super-resolution measurements were performed on a custom-made inverted microscope based on a Nikon Eclipse Ti-E frame with an oil immersion objective (Nikon CFI Apo TIRF 100XC Oil, NA = 1.49). Epifluorescence illumination was applied at an excitation wavelength of 647 nm. The laser power density was set to 2–4 kW/cm^2^ on the sample plane and controlled via an acousto-optic tunable filter. The separation of excitation from emission wavelengths was achieved by using a filter set from Semrock (Di03-R405/488/561/635-t1-25 × 36 BrightLine® quad-edge super-resolution/TIRF dichroic beam-splitter and FF01-446/523/600/677-25 BrightLine® quad-band bandpass filter, and an additional AHF 690/70 H emission filter). Images of individual fluorescent dye molecules were captured by an Andor iXon3 897 BV EMCCD camera (512 × 512 pixels with 16 μm pixel size) with the following acquisition parameters: exposure time = 20 ms; EM gain = 200; temperature = -75°C. For all presented images, 20,000 frames were captured from a single region of interest. The Nikon Perfect Focus System was used to keep the sample in focus during measurements. High-resolution images were reconstructed with the rainSTORM localization software^55^. The axial position of each individual fluorescent molecule was obtained by introducing astigmatism into the system via a cylindrical lens placed in front of the detector and measuring the ellipticity of the point spread function of each fluorescent molecule. This astigmatic 3D super-resolution technique^56^ was calibrated using 20 nm fluorescent polystyrene beads moved axially by 25 nm steps through the depth of field with a piezo stage. Spatial drift introduced by either mechanical movement or thermal effects was analyzed and reduced using an autocorrelation-based blind drift correction algorithm. The detected localizations were filtered based on the fitting parameters (e.g., sigma values, residue, intensity). The pixel size of the final pixelized super-resolution images was set at 20 nm, based on the localization precision and localization density. For junctional image analysis, we first determined the path along cell-cell junctions on super-resolved 2D localisation density images. To achieve this, two endpoints were manually marked on the junction, and the path between the end points along the junction was outlined similarly to Gray *et al*.^57^ by the algorithm, balancing the shortest path with the regions of highest density. The path was then smoothed and resampled using spline fitting. This marked path formed the basis for further analyses. To determine junctional width, the histogram of the signed distances of localisation points from the marked path were first obtained using a modified ‘distance2curve’ MATLAB function. Then, the autoconvolution of the histogram was calculated and the background was removed. The junctional width value was given as the full width at half maximum of the resulting peak. To make side view projections of tight junctions, claudin-5 localisations were projected onto the marked path in the x-y plane, while maintaining their axial (z) coordinates. The positions of projections along the path and the axial coordinates of localisations were then used as the new point coordinates for the side view. Zig-zaginess/jagedness of junctional segments was calculated as the relative difference of the marked path length to the Euclidean distance of the endpoints.

### Confocal microscopy and image analysis of *in vitro* samples

Confocal microscopy of *in vitro* samples was performed using Leica TCS SP5 AOBS and Leica Stellaris confocal laser scanning microscopes (Leica Microsystems) equipped with HC PL APO 20× (NA=0.7) and HCX PL APO 63× oil (NA=1.4) objectives. 3D renderings as well as x-z and y-z projections were generated in Leica Application Suite X software v3.7.5 (Leica Microsystems) on a series of z-stacks with 0.1 μm spacing. Image analysis was performed in FIJI/ImageJ on 8-bit, grayscale images. The staining intensity of focal adhesion proteins was determined using the ‘Mean gray value’ function. The calculation of cell ratios and subcellular clustering of focal adhesion staining was done manually in FIJI/ImageJ using a set of criteria described in **Supplementary Fig. 3a** and **Supplementary Fig. 6c**.

### Fluidic force microscopy (FluidFM)

A robotic fluidic force microscopy setup (FluidFM OMNIUM, Cytosurge AG) was used to measure direct adhesion forces of individual ECs within a monolayer during BBB maturation. Human stem cell-derived ECs were seeded in 6-well plates (Corning, cat# 3736) coated with a mixture of collagen IV (100 µg/mL) and fibronectin (25 µg/mL) at a seeding density of 2.5×10^4^ cells/cm^2^. Confluent monolayers were treated with cARLA or control medium for 48 h to induce BBB maturation. FluidFM measurements were performed in a 37 °C incubator with 5% CO_2_. Before measurements, a micropipette probe (8 µm aperture diameter; 2 N/m nominal spring constant; Cytosurge AG) was inserted into the device. The microfluidic channel inside the probe was filled with 1 µL of Dulbecco’s phosphate buffered saline (DPBS, Sigma-Aldrich). The probe was then mounted onto the FluidFM head, and the laser was manually aligned at the end of the cantilever. The laser signal reaches the position-sensitive detector reflecting from an automatically aligned mirror, which provides optimal light distribution between the sensor segments^58^. Inverse optical lever sensitivity was measured on the surface of a medium-filled well without cells. To convert to distances, values measured by the position-sensitive detector were multiplied by the sensitivity and the spring constant^59^. ECs within a monolayer were approached with a speed of 10 µm/s, and the setpoint defining the mechanical contact between the cell and surface was 200 nN. After reaching the setpoint, a -500 mbar negative pressure was applied in the microfluidic channel inside the cantilever. When the cantilever was in contact with ECs for 30 seconds, it was retracted to a height of 100 µm with a retraction speed of 1 µm/s. Force-distance curves were generated from measured values, which were used to analyse adhesion forces and the work of detachment. Adhesion forces were defined as force values at absolute minima of curves, while the work of detachment was defined as the area under the curve.

### Atomic force microscopy (AFM)

AFM experiments were performed using an Asylum Research MFP-3D atomic force microscope (Asylum Research; software in IgorPro 6.34A, Wavemetrics) mounted on top of a Zeiss Axiovert 200 optical microscope, which was used for optical positioning. Human stem cell-derived ECs were grown on 35 mm Petri dish lids (Greiner Bio-One, cat# 627102) coated with collagen IV (100 μg/mL) and fibronectin (25 μg/mL) at a seeding density of 2.5×10^4^ cells/cm^2^. Confluent monolayers were treated with cARLA or control medium for 48 h to induce BBB maturation. Measurements were performed using V-shaped rectangular overall gold-coated cantilevers (BioLever A-short lever; 30 pN/nm nominal spring constant; 37 kHz resonant frequency in air; Olympus). The spring constant of the cantilever was determined each time by thermal calibration^60^. To compare the elastic properties of ECs, 80 µm×80 µm areas were selected and divided into 40 lines by 40 columns. In each point, a single force curve was recorded with a trigger of 300 pN, total travelling distance of 2 µm, acquisition frequency of 2 Hz, resulting a travelling speed of 8 µm/s both for landing and retraction of the cantilever. From each recorded curve, Young’s modulus and elasticity index values were calculated and stitched together into a pseudo-colored elastic map in MATLAB as previously described^61^.

### Resonant waveguide grating (RWG)

Real-time kinetics of cell adhesion was measured using the Epic Benchtop System (Corning) instrument, which features a high-resolution, high-throughput optical sensor based on RWG technology. Human stem cell-derived ECs were cultured in specialised, fibronectin-coated 384-well microplates (Corning, cat# 5040), in which each well contains a 2×2 mm² RWG sensor area, at a seeding density of 2.5×10^4^ cells/cm^2^. The microplate is illuminated from below by a tunable broadband light source with 825-840 nm wavelengths. Light is coupled through the grating areas into a thin waveguide layer with high-refractive-index made from biocompatible Nb₂O₅, which is deposited on a glass substrate. Within the waveguide, the coupled light undergoes multiple total internal reflections before being detected by a CCD camera. This propagation generates an evanescent electromagnetic field, which decays exponentially with distance from the sensor surface, allowing interaction with the sample. The evanescent field extends into the suspension above the surface, penetrating a 150 nm-thick region in contact with the waveguide. Using this system, wavelength shifts – which are directly proportional to the surface area covered by and local density of cell-matrix adhesions – are detected with a sensitivity of 0.25 pm^24^. After ECs reached confluency in wells, the baseline was recorded for 5 minutes. EC monolayers were then treated with cARLA or control medium using an electronic 16-channel pipette. Measurements were performed for 90-120 min at room temperature immediately after treatment, 24 h after treatment and 48 h after treatment. Between measurements, ECs were returned to the 37 °C incubator with 5% CO_2_. To investigate the effect of trypsin on cell-cell and cell-matrix adhesions, ECs were treated with 0.05% trypsin-EDTA for an additional 10 min after the 48 h measurement. Resonant wavelength shifts were calculated and compared to both the baseline and relative to the control group.

### Generation of *Cldn5* overexpressing and knockout bEnd.3 cells

The mouse immortalised brain endothelial cell line bEnd.3 was cultured in Dulbecco’s modified Eagle’s medium (DMEM; Sigma-Aldrich, cat# D5030) supplemented with 10% FBS (Sigma-Aldrich, cat# F4135), 100 U/mL penicillin, and 100 μg/ml streptomycin (Sigma-Aldrich, cat# P4333) in a humidified incubator at 37 °C with 5% CO_2_. Mouse bEnd.3 cells overexpressing human CLDN-5 (CLDN5 OE) or lacking mouse CLDN-5 (Cldn5 KO) were generated as described previously^62^. Briefly, for CLDN5 OE cells, pcDNA3.1 (+) encoding human CLDN-5 was transfected into cells using FuGENE HP DNA (Promega) and cells were cultured for more than 10 days with 10 μg/mL blasticidin S (Invivogen). For Cldn5 KO cells, Cas9 was transduced into cells by LentiArray Cas9 Lentivirus (Thermo Fisher, cat# A32064) and gRNA against mouse *Cldn5* (target sequence: 5’-CCGTCGGATCATAGAACTCG-3’) expressing pU6 vector with puromycin resistance gene was transfected into cells. Cells were then cultured for 2 weeks in medium supplemented with 3 μg/mL puromycin (Invivogen) and 10 μg/mL blasticidin S. A limiting dilution was performed to isolate single colony-derived cells.

### RNA-sequencing and analysis

RNA was isolated from confluent monolayers of *Cldn5* OE, KO and mock-transfected bEnd.3 cells using the E.Z.N.A. Total RNA Kit (Omega Bio-tek, cat# R6834) according to the manufacturer’s instructions. Residual DNA was digested by DNase I on column, and cDNA was prepared using the High Capacity cDNA Reverse Transcription Kit (Applied Biosystems, cat# 4368814). RNA-sequencing and data processing were performed at Macrogen Europe BV (Amsterdam, The Netherlands) using the TruSeq Stranded mRNA LT Sample Prep Kit, 100 bp paired-end reads and 20 million reads per sample, on a NovaSeq 6000 instrument (Illumina). Raw FASTQ files were trimmed with Cutadapt and aligned to the mouse reference genome (version GRCm39) using STAR^63^. The resulting BAM files were sorted with STAR and indexed with Samtools. Gene quantification was performed using the RSEM’s rsem-calculate-expression tool. The following published datasets were also reanalysed in this study: control and cARLA-treated human stem cell-derived ECs^22^; GEO accession number: GSE224846) as well as FACS-isolated mouse brain ECs in acute epilepsy vs. control^26^; GEO accession number: GSE111839). Differential expression analysis was performed using the DESeq2 R/Bioconductor package, in which log_2_(fold change) and *P*-values were obtained for each gene. To account for multiple comparisons, false discovery rate (FDR) was calculated using the Benjamini-Hochberg method. Genes with an FDR < 0.01 and log_2_FC > 0.3 or log_2_FC < -0.3 were considered to be differentially expressed. The g:GOSt tool in g:Profiler^64^ was used to identify over-represented Gene Ontology (GO) terms in acute epilepsy and to calculate gene ratios and statistical significance for each pathway. Heatmaps were created using Min-Max scaling on transcript per million (TPM) normalized counts in Prism 10 (GraphPad).

### Immunohistochemistry and image analysis of *ex vivo* samples

Mice were sacrificed by cervical dislocation and brains were frozen in O.C.T. reagent (Fisher Scientific, cat# 23-730-571). Resected brain tissue from TLE patients was rapidly frozen on dry ice, embedded in O.C.T. reagent, and 20 μm sections from the lateral temporal lobe and hippocampus were prepared. TLE and autopsy control sections were fixed in ice cold methanol for 10 min, and blocked in 5% BSA in PBS containing 0.2% Triton X-100 for 30 min at room temperature. Incubation with primary antibodies (in blocking buffer) were performed overnight at 4 °C. Sections were then incubated with secondary antibodies in PBS containing 0.2% Triton X-100 for 1 h at room temperature. Antibodies used are listed in **Supplementary Table 4**. Between each step, slides were washed in PBS. Slides were coverslipped with Aqua-Poly/Mount (Polysciences, cat# 18606-5). Images were acquired on a Zeiss Apotome 3 optical sectioning microscope equipped with an Axiocam 705 mono sCMOS camera and Colibri 5 LED illuminator with 385nm/475nm/555nm/630nm excitation using Zen Blue software v3.8.3. Image analysis was performed in FIJI/ImageJ on 8-bit, grayscale images. Blood vessels as regions of interest were outlined and masked by automatic thresholding with manual adjustments defined by vascular (collagen IV) and tight junction (claudin-5) stainings. Claudin-5 vessel area fraction was determined using the ‘Area fraction’ function within regions of interest. Zyxin and vinculin intensities were quantified within regions of interest (vascular) or outside these regions (parenchymal) using the ‘Mean gray value’ function.

### Statistics

No statistical method was used to pre-determine sample sizes. All key experiments were repeated independently, and detailed information about error bars, sample size, definition of replicates, statistical tests and *P*-values are provided in each figure legend. Statistical analyses were performed using Prism 10 (GraphPad). Normality was determined by D’Agostino-Pearson, Shapiro-Wilk and Kolmogorov-Smirnov normality tests. On normally distributed data, unpaired t-test (two-tailed) or one-way ANOVA followed by Dunnett’s post-hoc test were used to compare means from two or more groups, respectively. If standard deviations were different between groups (determined by an F-test), Welch’s correction was applied. On non-normally distributed data, a non-parametric Mann-Whitney test (two-tailed) was used to compare means from two groups, and relationships between two variables were determined using a Spearman correlation. To account for multiple comparisons in RNA-sequencing data, false discovery rate (FDR) was calculated using the Benjamini-Hochberg method. Statistical significance was generally set at P < 0.05, and FDR < 0.01 for RNA-sequencing analysis.

## Data availability

RNA-seq data generated/analysed in this study can be found in the Gene Expression Omnibus (GEO) repository under accession numbers GSE307207, GSE224846 and GSE111839. All other data supporting the findings from this study are available within the manuscript and the Supplementary information. Source data are provided with this manuscript.

## Acknowledgements

We thank the support of the Cellular Imaging Laboratory (HUN-REN BRC) and Dr. Ana Martins for the critical reading of our manuscript.

M.A.D. was funded by the National Research, Development and Innovation Office of Hungary (K143766; 2024-1.2.2-ERA_NET-2024-00018) and the Hungarian Academy of Sciences (NAP2022-I-6/2022). R.H. was funded by the TKP2021 funding scheme (TKP2021-EGA04) and KDP-2021 programs provided by the Ministry for Innovation and Technology of Hungary from the National Research, Development and Innovation Fund; the HUN-REN Hungarian Research Network and by the Lendület Program of the Hungarian Academy of Sciences. M.Ca. was funded by Taighde Éireann – Research Ireland, (Eye-D-21/SPP/3732 and 21/RC/10294_P2 at FutureNeuro Research Ireland Centre for Translational Brain Science), and was supported by the European Research Council (ERC – Retina-Rhythm, 864522). D.H. was funded by Research Ireland grants 16/RC/3948 and 21/RC/10294_P2 (FutureNeuro). M.E. was funded by 2022-2.1.1-NL-2022-00012 provided by the Ministry of Culture and Innovation of Hungary from the National Research, Development and Innovation Fund, financed under the 2022-2.1.1-NL funding scheme, and TKP2021-NVA-19 provided by the Ministry of Culture and Innovation of Hungary from the National Research, Development and Innovation Fund, financed under the TKP2021-NVA funding scheme. G.P. was supported by the National Academy of Scientist Education Program of the National Biomedical Foundation under the sponsorship of the Hungarian Ministry of Culture and Innovation; and the the European Union’s Horizon 2020 research and innovation programme under the Marie Skłodowska-Curie grant agreement (101034252). I.R. was supported by the Gedeon Richter Talentum Foundation in framework of Gedeon Richter Excellence PhD Scholarship of Gedeon Richter. E.F. was supported by the STARTING_24 program (STARTING 150012) of by the National Research, Development and Innovation Office (NKFIH) in Hungary. C.G. was supported by start-up funding from the StAR programme at RCSI and a CURE Epilepsy Taking Flight Award. The funders had no role in study design, data collection and analysis, decision to publish or preparation of the manuscript.

## Author contributions

G.P., M.Ca., R.H. and M.A.D. designed research; G.P., L.L., B.M., I.R., T.N., B.H.K., I.G., A.S., K.D.K., E.F., B.K., I.S., A.G.V., Y.H., C.G., A.M., S.V. performed experiments and analysed data; M.Cu., D.C.H., K.J.S., D.F.OB., M.E., contributed reagents/analytic tools; M.Ca., R.H. and M.A.D. jointly supervised the study; G.P., M.Ca., R.H. and M.A.D. wrote the paper, which was edited by all authors.

## References

1. Abbott, N.J. & Friedman, A. Overview and introduction: The blood–brain barrier in health and disease. Epilepsia 53, 1–6 (2012).

2. Aitken, C., Mehta, V., Schwartz, M.A. & Tzima, E. Mechanisms of endothelial flow sensing. Nat Cardiovasc Res 2, 517–529 (2023).

3. Hansen, C.E., Hollaus, D., Kamermans, A. & de Vries, H.E. Tension at the gate: sensing mechanical forces at the blood–brain barrier in health and disease. Journal of Neuroinflammation 21, 325 (2024).

4. Konig, S., Jayarajan, V., Wray, S., Kamm, R. & Moeendarbary, E. Mechanobiology of the blood-brain barrier during development, disease and ageing. Nature Communications 16, 7233 (2025).

5. Harraz, O.F., Klug, N.R., Senatore, A.J., Hill-Eubanks, D.C. & Nelson, M.T. Piezo1 Is a Mechanosensor Channel in Central Nervous System Capillaries. Circulation Research 130, 1531–1546 (2022).

6. Lim, X.R. et al. Endothelial Piezo1 channel mediates mechano-feedback control of brain blood flow. Nature Communications 15, 8686 (2024).

7. Hansen, C.E. et al. Inflammation-induced TRPV4 channels exacerbate blood–brain barrier dysfunction in multiple sclerosis. Journal of Neuroinflammation 21, 72 (2024).

8. Fu, B.M. & Tarbell, J.M. Mechano-sensing and transduction by endothelial surface glycocalyx: composition, structure, and function. WIREs Systems Biology and Medicine 5, 381–390 (2013).

9. Ben-Zvi, A. et al. Mfsd2a is critical for the formation and function of the blood–brain barrier. Nature 509, 507–511 (2014).

10. Andreone, B.J. et al. Blood-Brain Barrier Permeability Is Regulated by Lipid Transport-Dependent Suppression of Caveolae-Mediated Transcytosis. Neuron 94, 581–594.e585 (2017).

11. Berndt, P. et al. Tight junction proteins at the blood-brain barrier: far more than claudin-5. Cell Mol Life Sci 76, 1987–2002 (2019).

12. Greene, C., Hanley, N. & Campbell, M. Claudin-5: gatekeeper of neurological function. Fluids and Barriers of the CNS 16, 3 (2019).

13. Hashimoto, Y., Greene, C., Munnich, A. & Campbell, M. The CLDN5 gene at the blood-brain barrier in health and disease. Fluids and Barriers of the CNS 20, 22 (2023).

14. van Vliet, E.A. et al. Blood–brain barrier leakage may lead to progression of temporal lobe epilepsy. Brain 130, 521–534 (2006).

15. Marchi, N. et al. Seizure-promoting effect of blood-brain barrier disruption. Epilepsia 48, 732–742 (2007).

16. Friedman, A. Blood-brain barrier dysfunction, status epilepticus, seizures, and epilepsy: a puzzle of a chicken and egg? Epilepsia 52 Suppl 8, 19–20 (2011).

17. Rüber, T. et al. Evidence for peri-ictal blood–brain barrier dysfunction in patients with epilepsy. Brain 141, 2952–2965 (2018).

18. Rempe, R.G. et al. Matrix Metalloproteinase-Mediated Blood-Brain Barrier Dysfunction in Epilepsy. The Journal of Neuroscience 38, 4301–4315 (2018).

19. Greene, C. et al. Microvascular stabilization via blood-brain barrier regulation prevents seizure activity. Nature Communications 13, 2003 (2022).

20. Reiss, Y. et al. The neurovasculature as a target in temporal lobe epilepsy. Brain Pathology 33, e13147 (2023).

21. Martin, M. et al. Engineered Wnt ligands enable blood-brain barrier repair in neurological disorders. Science 375, eabm4459 (2022).

22. Porkoláb, G. et al. Synergistic induction of blood–brain barrier properties. Proceedings of the National Academy of Sciences 121, e2316006121 (2024).

23. Nagy, Á.G., Székács, I., Bonyár, A. & Horvath, R. Cell-substratum and cell-cell adhesion forces and single-cell mechanical properties in mono- and multilayer assemblies from robotic fluidic force microscopy. European Journal of Cell Biology 101, 151273 (2022).

24. Orgovan, N. et al. Dependence of cancer cell adhesion kinetics on integrin ligand surface density measured by a high-throughput label-free resonant waveguide grating biosensor. Scientific Reports 4, 4034 (2014).

25. Greene, C. et al. Dose-dependent expression of claudin-5 is a modifying factor in schizophrenia. Molecular Psychiatry 23, 2156–2166 (2018).

26. Munji, R.N. et al. Profiling the mouse brain endothelial transcriptome in health and disease models reveals a core blood–brain barrier dysfunction module. Nature Neuroscience 22, 1892–1902 (2019).

27. Sasson, E. et al. Nano-scale architecture of blood-brain barrier tight-junctions. eLife 10, e63253 (2021).

28. Bell, B., Anzi, S., Sasson, E. & Ben-Zvi, A. Unique features of the arterial blood–brain barrier. Fluids and Barriers of the CNS 20, 51 (2023).

29. Tani, E., Yamagata, S. & Ito, Y. Freeze-fracture of capillary endothelium in rat brain. Cell and Tissue Research 176, 157–165 (1977).

30. Shivers, R.R., Betz, A.L. & Goldstein, G.W. Isolated rat brain capillarie spossess intact, structurally complex, interendothelial tight junctions; freeze-fracture verification of tight junction integrity. Brain Research 324, 313–322 (1984).

31. Wolburg, H. et al. Modulation of tight junction structure in blood-brain barrier endothelial cells. Effects of tissue culture, second messengers and cocultured astrocytes. J Cell Sci 107 (Pt 5), 1347–1357 (1994).

32. Liebner, S., Kniesel, U., Kalbacher, H. & Wolburg, H. Correlation of tight junction morphology with the expression of tight junction proteins in blood-brain barrier endothelial cells. European Journal of Cell Biology 79, 707–717 (2000).

33. Zihni, C., Mills, C., Matter, K. & Balda, M.S. Tight junctions: from simple barriers to multifunctional molecular gates. Nature Reviews Molecular Cell Biology 17, 564–580 (2016).

34. Gonschior, H. et al. Nanoscale segregation of channel and barrier claudins enables paracellular ion flux. Nature Communications 13, 4985 (2022).

35. Khatau, S.B. et al. A perinuclear actin cap regulates nuclear shape. Proceedings of the National Academy of Sciences 106, 19017–19022 (2009).

36. Kim, D.-H. et al. Actin cap associated focal adhesions and their distinct role in cellular mechanosensing. Scientific Reports 2, 555 (2012).

37. Chambliss, A.B. et al. The LINC-anchored actin cap connects the extracellular milieu to the nucleus for ultrafast mechanotransduction. Scientific Reports 3, 1087 (2013).

38. Schoofs, H. et al. Dynamic cytoskeletal regulation of cell shape supports resilience of lymphatic endothelium. Nature 641, 465–475 (2025).

39. Végh, A.G. et al. Spatial and temporal dependence of the cerebral endothelial cells elasticity. Journal of Molecular Recognition 24, 422–428 (2011).

40. Kataoka, N. et al. Measurements of endothelial cell-to-cell and cell-to-substrate gaps and micromechanical properties of endothelial cells during monocyte adhesion. Proceedings of the National Academy of Sciences 99, 15638–15643 (2002).

41. Pesen, D. & Hoh, J.H. Micromechanical Architecture of the Endothelial Cell Cortex. Biophysical Journal 88, 670–679 (2005).

42. Mehta, D. et al. Modulatory role of focal adhesion kinase in regulating human pulmonary arterial endothelial barrier function. The Journal of Physiology 539, 779–789 (2002).

43. Wu, M.H. Endothelial focal adhesions and barrier function. J Physiol 569, 359–366 (2005).

44. Ayloo, S. et al. Pericyte-to-endothelial cell signaling via vitronectin-integrin regulates blood-CNS barrier. Neuron 110, 1641–1655.e1646 (2022).

45. Arnold, K.M., Goeckeler, Z.M. & Wysolmerski, R.B. Loss of Focal Adhesion Kinase Enhances Endothelial Barrier Function and Increases Focal Adhesions. Microcirculation 20, 637–649 (2013).

46. Jean, C. et al. Inhibition of endothelial FAK activity prevents tumor metastasis by enhancing barrier function. Journal of Cell Biology 204, 247–263 (2014).

47. Lee, J., Borboa, A.K., Chun, H.B., Baird, A. & Eliceiri, B.P. Conditional deletion of the focal adhesion kinase FAK alters remodeling of the blood-brain barrier in glioma. Cancer Res 70, 10131–10140 (2010).

48. Kwan, P., Schachter, S.C. & Brodie, M.J. Drug-resistant epilepsy. N Engl J Med 365, 919–926 (2011).

49. Garcia, F.J. et al. Single-cell dissection of the human brain vasculature. Nature 603, 893–899 (2022).

50. Yang, A.C. et al. A human brain vascular atlas reveals diverse mediators of Alzheimer’s risk. Nature 603, 885–892 (2022).

51. Winkler, E.A. et al. A single-cell atlas of the normal and malformed human brain vasculature. Science 375, eabi7377 (2022).

52. Wälchli, T. et al. Single-cell atlas of the human brain vasculature across development, adulthood and disease. Nature 632, 603–613 (2024).

53. Cecchelli, R. et al. A stable and reproducible human blood-brain barrier model derived from hematopoietic stem cells. PLoS One 9, e99733 (2014).

54. van de Linde, S. et al. Direct stochastic optical reconstruction microscopy with standard fluorescent probes. Nature Protocols 6, 991–1009 (2011).

55. Rees, E.J., Erdelyi, M., Schierle, G.S.K., Knight, A. & Kaminski, C.F. Elements of image processing in localization microscopy. Journal of Optics 15, 094012 (2013).

56. Huang, B., Wang, W., Bates, M. & Zhuang, X. Three-Dimensional Super-Resolution Imaging by Stochastic Optical Reconstruction Microscopy. Science 319, 810–813 (2008).

57. Gray, K.M., Katz, D.B., Brown, E.G. & Stroka, K.M. Quantitative Phenotyping of Cell-Cell Junctions to Evaluate ZO-1 Presentation in Brain Endothelial Cells. Ann Biomed Eng 47, 1675–1687 (2019).

58. Sztilkovics, M. et al. Single-cell adhesion force kinetics of cell populations from combined label-free optical biosensor and robotic fluidic force microscopy. Scientific Reports 10, 61 (2020).

59. Nagy, Á.G., Kámán, J., Horváth, R. & Bonyár, A. Spring constant and sensitivity calibration of FluidFM micropipette cantilevers for force spectroscopy measurements. Scientific Reports 9, 10287 (2019).

60. Sader, J.E. et al. Spring constant calibration of atomic force microscope cantilevers of arbitrary shape. Review of Scientific Instruments 83 (2012).

61. Varga, B. et al. Elasto-mechanical properties of living cells. Biochemistry and Biophysics Reports 7, 303–308 (2016).

62. Hashimoto, Y. et al. Recurrent de novo mutations in CLDN5 induce an anion-selective blood–brain barrier and alternating hemiplegia. Brain 145, 3374–3382 (2022).

63. Dobin, A. et al. STAR: ultrafast universal RNA-seq aligner. Bioinformatics 29, 15–21 (2013).

64. Kolberg, L. et al. g:Profiler-interoperable web service for functional enrichment analysis and gene identifier mapping (2023 update). Nucleic Acids Res 51 (2023).

