## Supplementary Figures 1-10, Supplementary Tables 1-4 for "Endothelial tight junctions and cell-matrix adhesions reciprocally control blood-brain barrier integrity"

**This file contains:**

Supplementary Figs. 1-10.

Supplementary Tables 1-4.

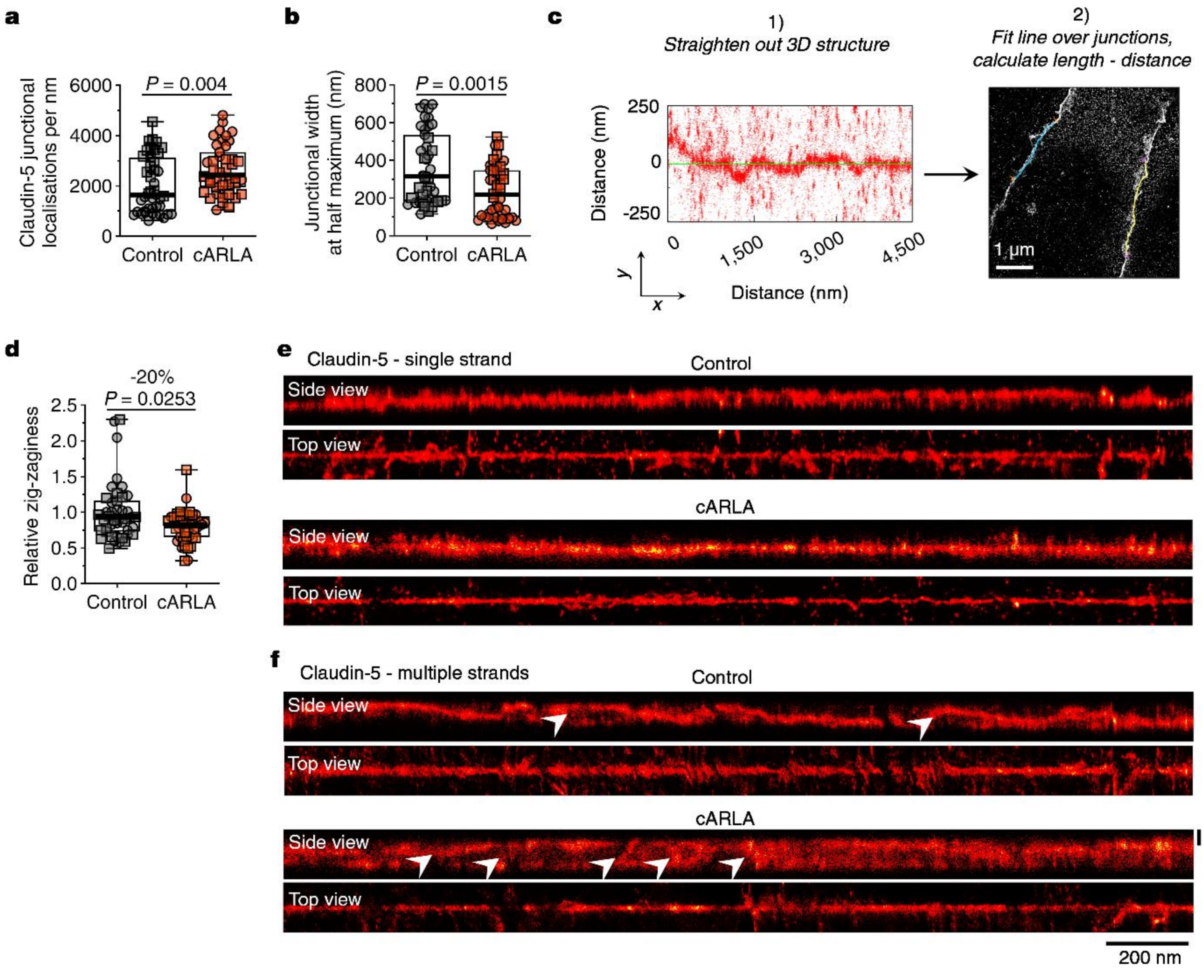


**Supplementary Fig. 1. Super-resolution imaging of claudin-5 in human stem cell-derived brain ECs. a)** Junctional claudin-5 localisations per nm. **b)** Tight junction width upon BBB maturation. Full width at half-maximum (FWHM) values derived from total claudin-5 distributions are shown. In both panels, box: median ± quartiles, whiskers: range. Mann-Whitney test, two-tailed, n=43 full-length junctions from two experiments. **c)** Method to determine zig-zaginess/jaggedness of tight junctions. **d)** Zig-zaginess of tight junctions relative to the control group. Box: median ± quartiles, whiskers: range. Mann-Whitney test, two-tailed, n=43 full-length junctions from two experiments. **e)** Representative super-resolution images of claudin-5$^+$ tight junctions composed of a single strand and **f)** composed of multiple strands in control and cARLA-treated human stem cell-derived brain ECs. Side (y/z) and top (x/y) projections from 3D dSTORM images are shown. White arrowheads indicate kissing points between tight junction strands.

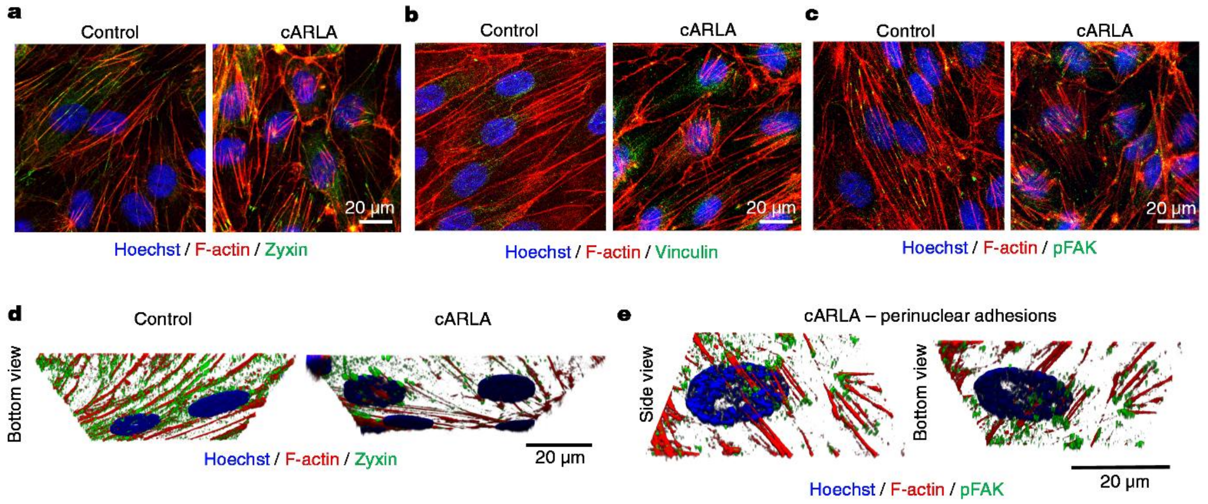


**Supplementary Fig. 2. Perinuclear adhesions are specific to the mature BBB and feature focal adhesion proteins. a)** Representative confocal microscopy images of F-actin co-stained with zyxin, **b)** vinculin and **c)** an active, phosphorylated form of the focal adhesion kinase (pFAK) in human stem cell-derived brain ECs. Nuclei were counterstained with Hoechst 33342. **d)** Representative 3D reconstructions of images show the unifom (control) and perinuclear (cARLA-treated) distribution of zyxin in ECs from below. **e)** Representative 3D reconstruction of cARLA-treated ECs show perinuclear adhesions containing pFAK.

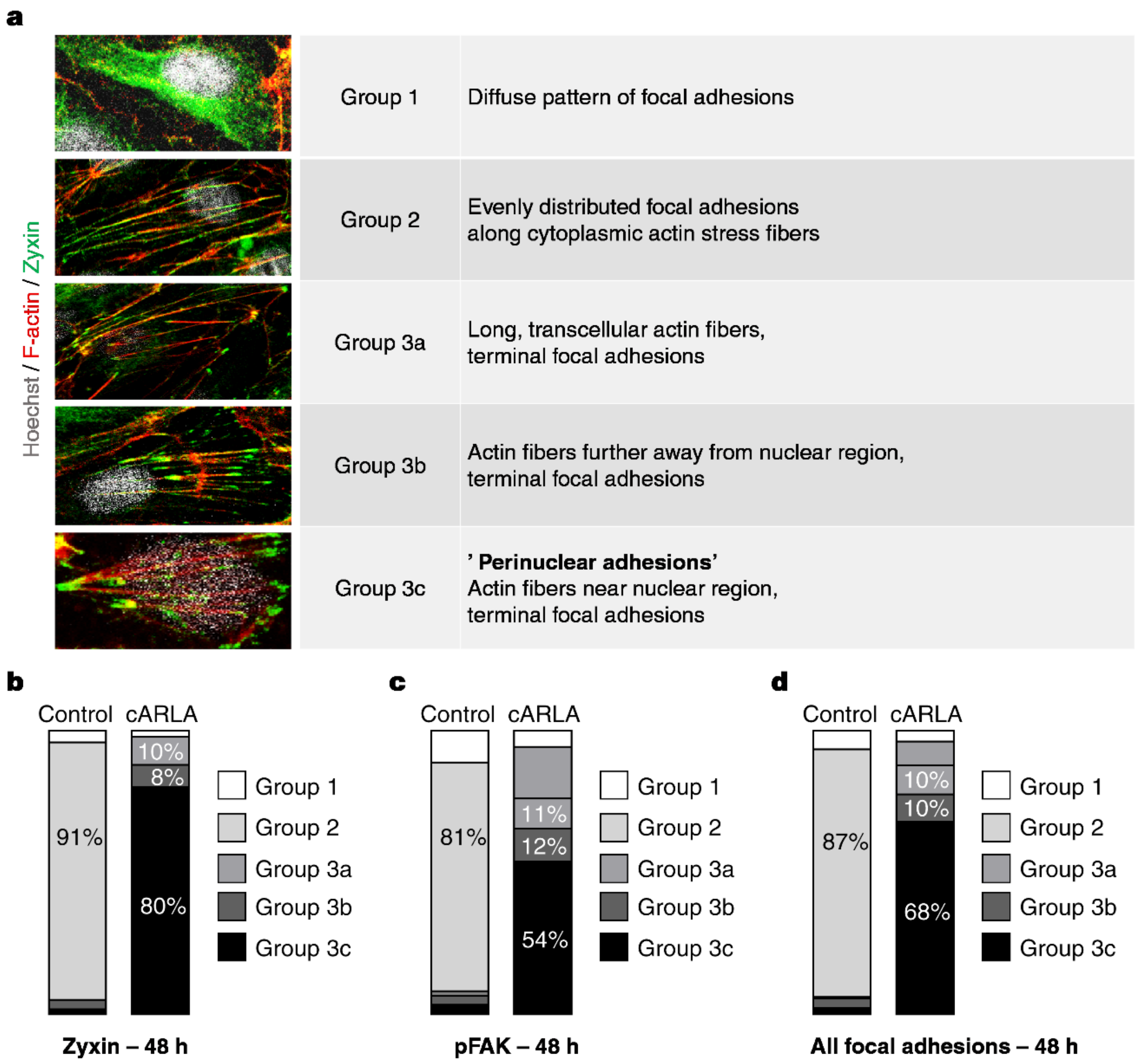


**Supplementary Fig. 3. Comprehensive distribution profile of focal adhesion proteins during BBB maturation. a)** Based on their staining pattern and subcellular localisation, 5 clusters of focal adhesion distributions were identified in human stem cell-derived brain ECs. **b)** Stacked bar plot showing the relative abundance of clusters during BBB maturation for zyxin, **c)** pFAK and **d)** combined. Note the marked increase in the abundance of Group 3c (perinuclear adhesions) in cARLA-treated ECs compared to the control group.

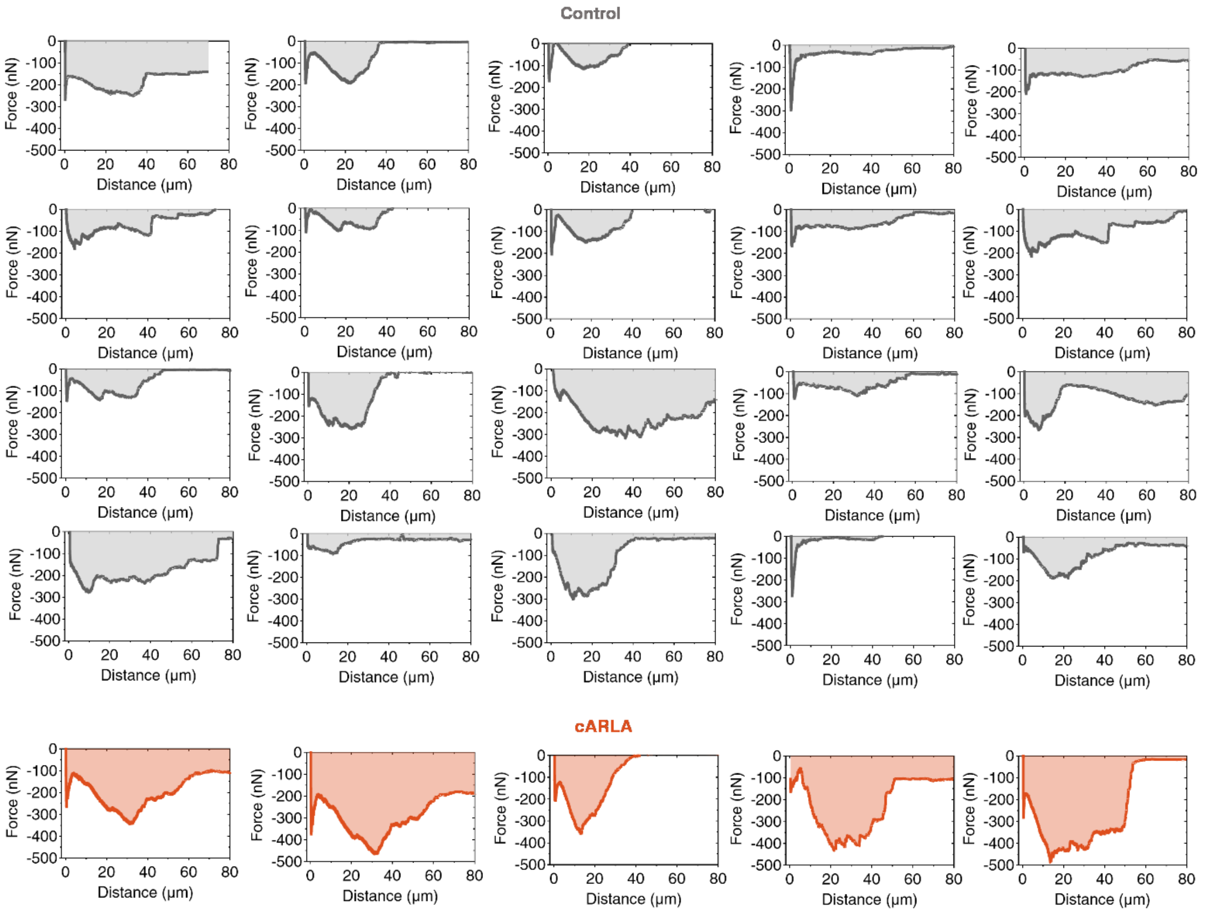


**Supplementary Fig. 4. Force-distance curves generated from FluidFM measurements.** This graph includes all successful cell detachment events from human stem cell-derived EC monolayers. Note the characteristic 2-dip distribution in all cARLA-treated ECs and most control ECs. The first dip corresponds to the disruption of cell-matrix adhesions, while the second, larger dip corresponds to the disruption of cell-cell junctions.

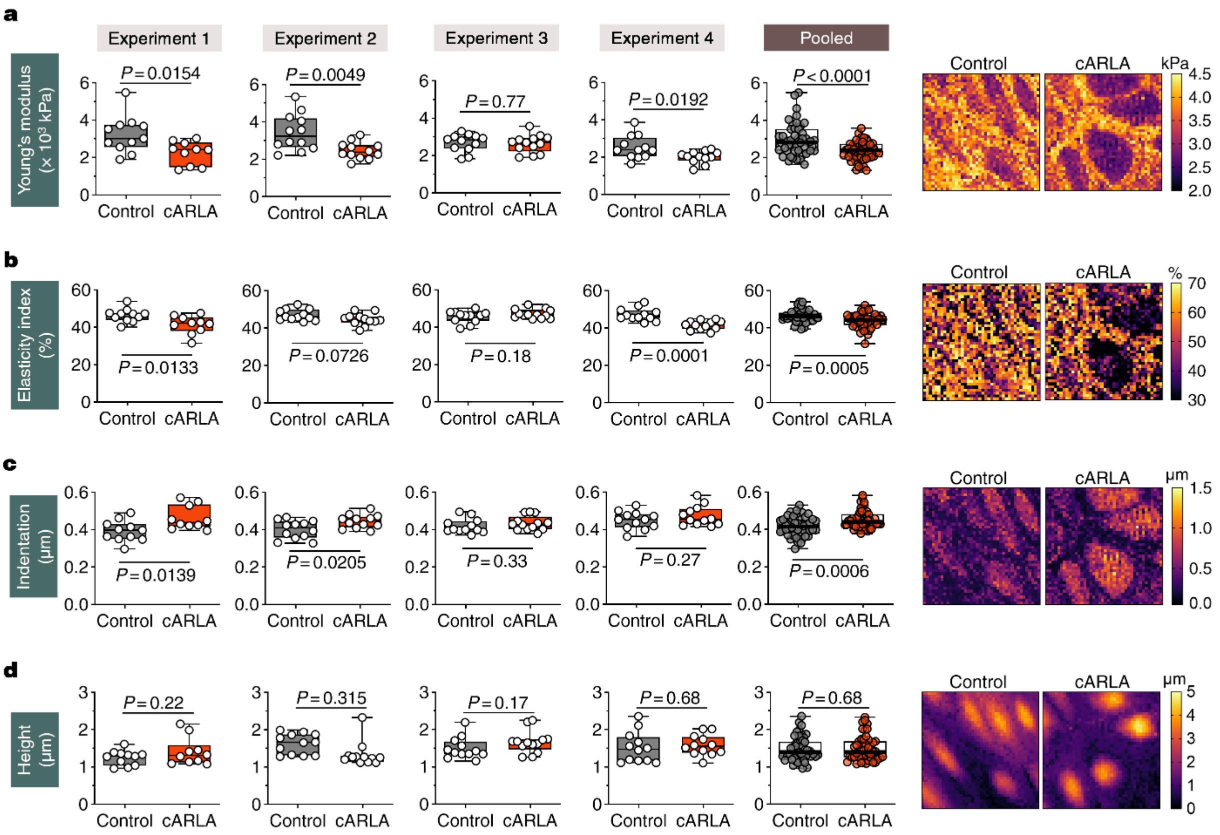


**Supplementary Fig. 5. Atomic force microscopy (AFM) measurements of BBB maturation in human stem cell-derived brain ECs. a)** Measured values for Young's modulus (overall hardness), **b)** elasticity index (higher values indicate an overall more elastic vs. plastic response to indentation with the probe), **c)** indentation and **d)** cell height. All values reflect an average derived from a 40 × 40µm area and do not differentiate between subcellular compartments. Colour-coded representative images (right panel) show differences between subcellular compartments. Experiments were repeated 4 times and values are shown both separately and pooled. In all panels, box: median ± quartiles, whiskers: range, unpaired t-test, two-tailed, n=47 images per group from 4 experiments.

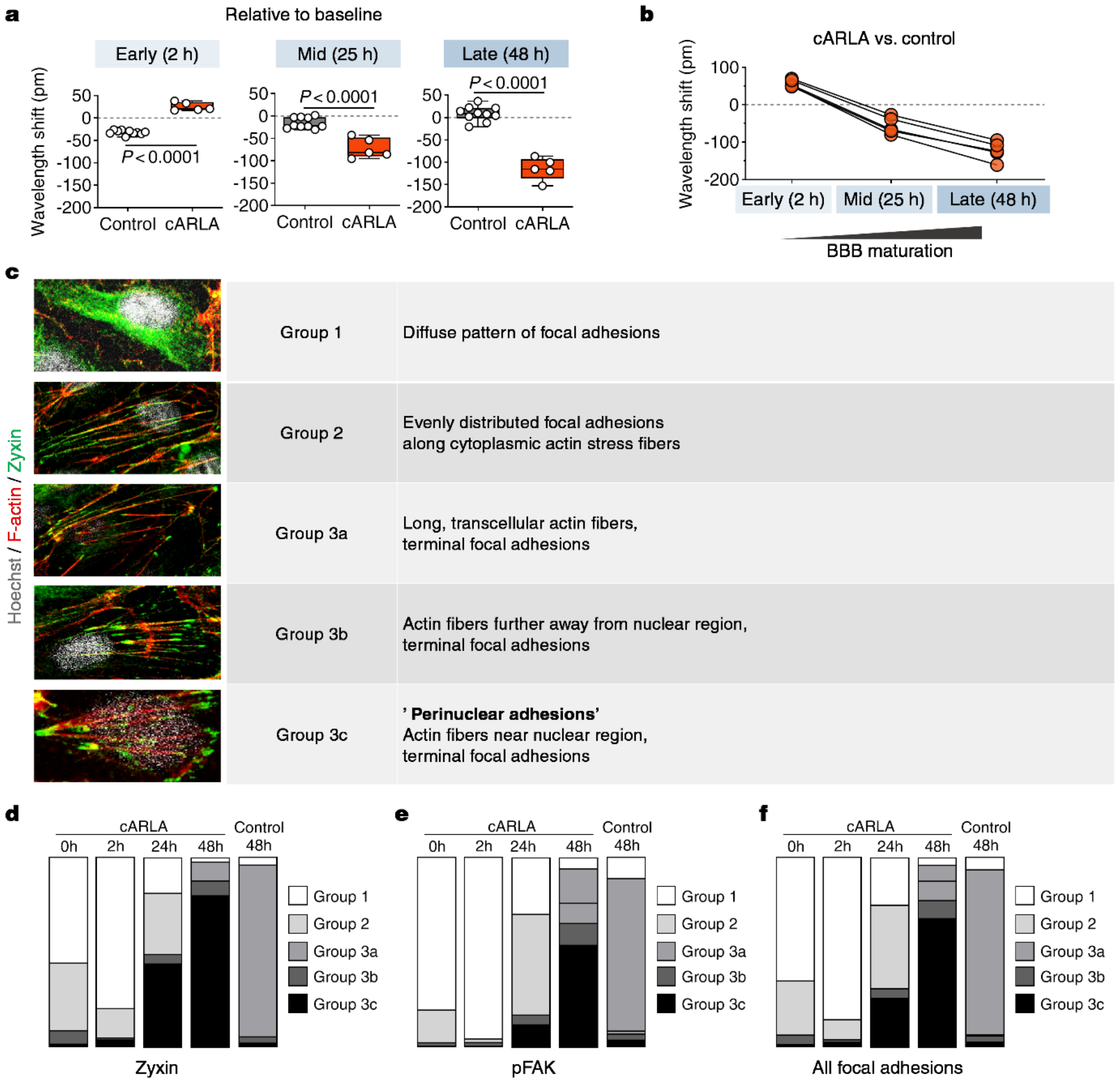


**Supplementary Fig. 6. Kinetics of cell-matrix adhesion and focal adhesion distribution during BBB maturation. a)** RWG wavelength shift values of human stem cell-derived brain ECs at early, mid and late timepoints of *in vitro* BBB maturation relative to baseline values and **b)** cARLA-treated ECs compared to control ECs. Note the progressive negative wavelength shift of cARLA-treated mature ECs during BBB maturation. **c)** Five clusters of focal adhesion distributions in ECs. **d)** Stacked bar plot showing the relative abundance of clusters over time during BBB maturation for zyxin, **e)** pFAK and **f)** combined. Note that the emergence of Group 3c (perinuclear adhesions) in cARLA-treated ECs starts at the mid timepoint (24 h) and is most prominent at the late timepoint (48 h) of barrier maturation.

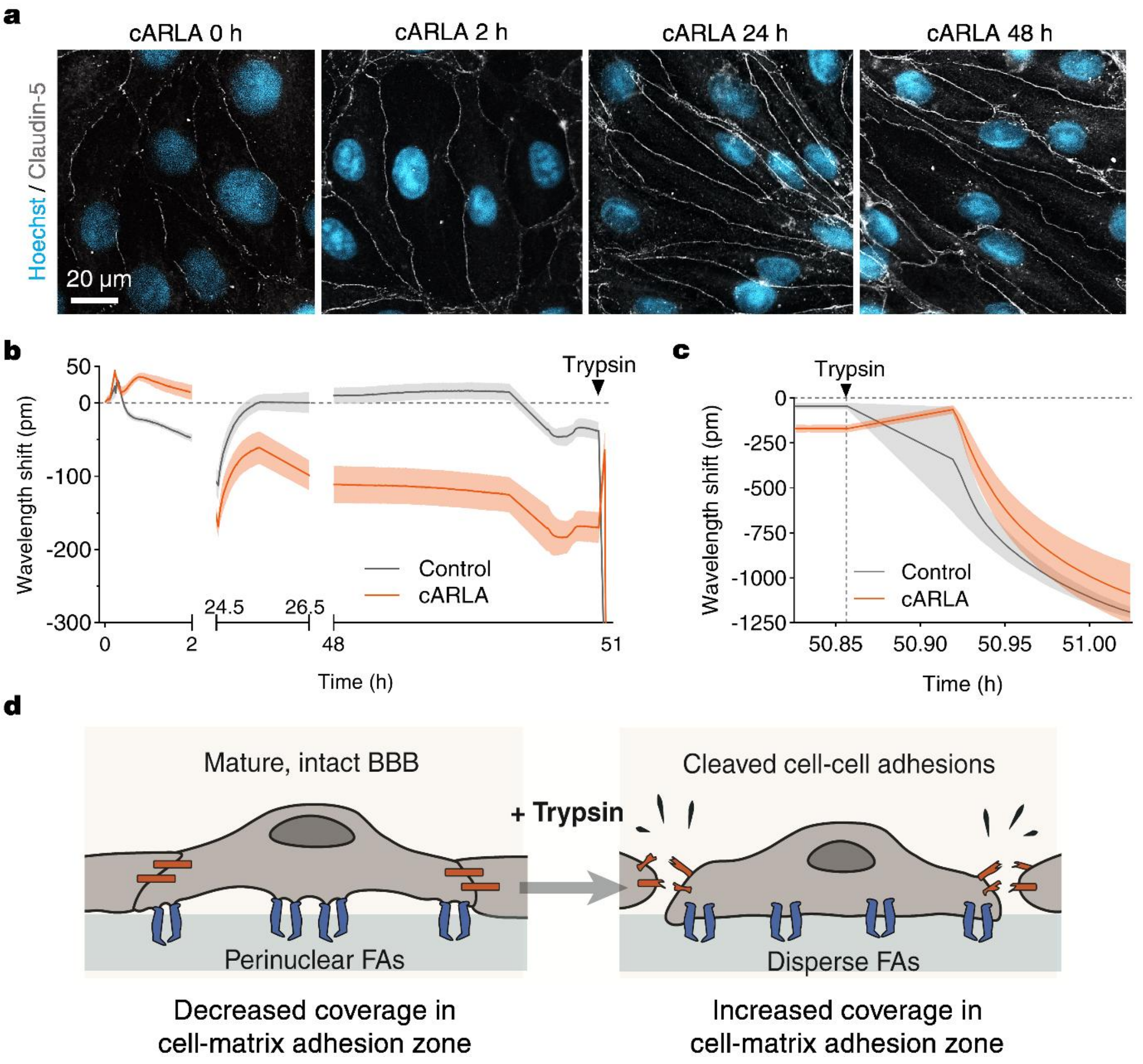


**Supplementary Fig. 7. Changes in cell-cell junctions precede changes in cell-matrix adhesions during BBB maturation. a)** Claudin-5 immunostaining in human stem cell-derived brain ECs treated with cARLA over time. Note the increasing continuity of claudin-5 staining at cell-cell junctions starting from 2 h. **b)** RWG wavelength shift kinetics of control and cARLA-treated mature ECs for the full timescale, and **c)** upon the addition of trypsin. Note the rapid positive wavelength shift in cARLA-treated ECs as trypsin cleaves cell-cell adhesions first, which has the same effect size as the effect of cARLA treatment vs. control beforehand. **d)** Shematic of the proposed mechanism of cell-cell junctions controlling the state of focal adhesions upon the addition of trypsin. FA: focal adhesion.

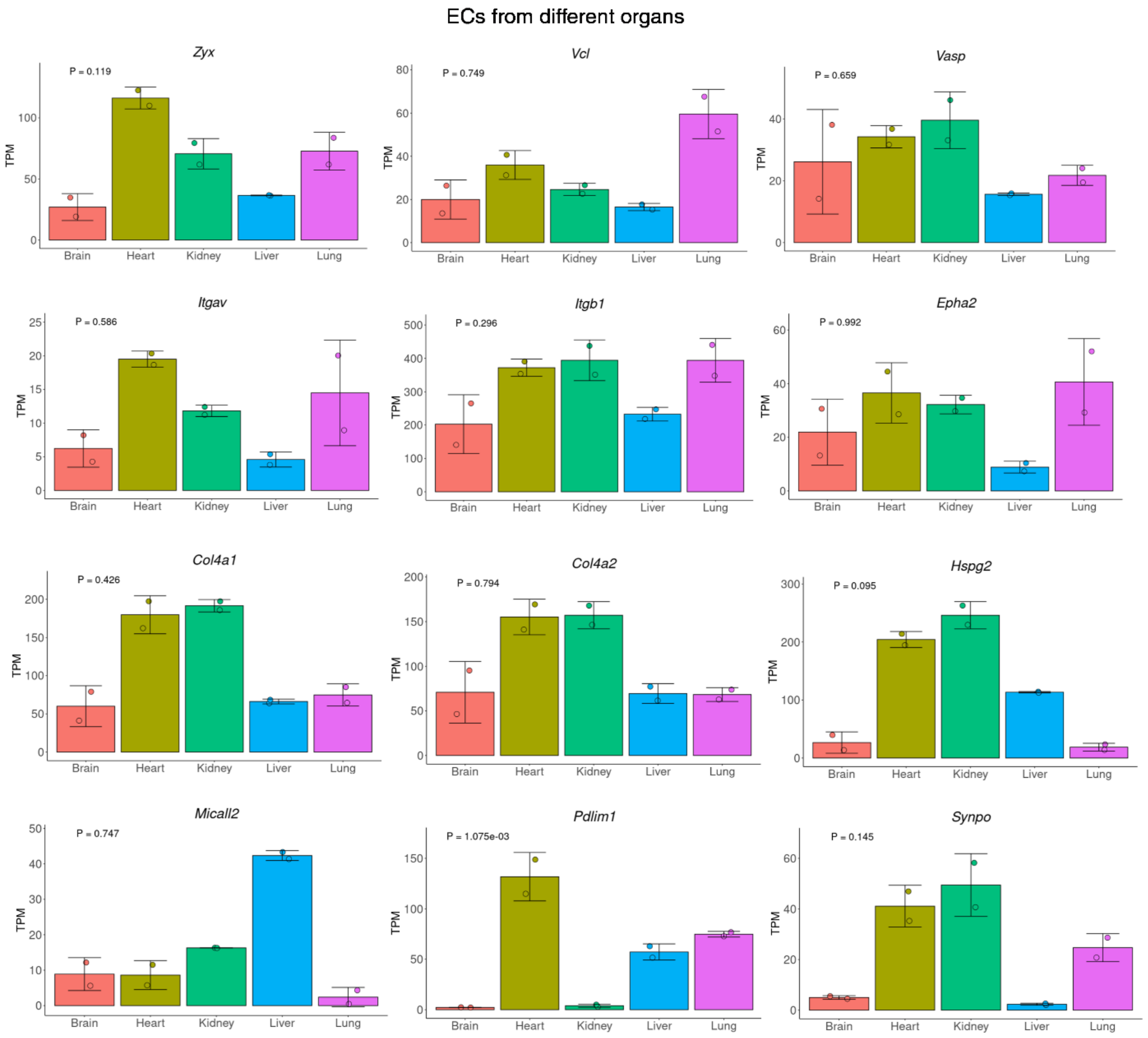


**Supplementary Fig. 8. Members of the core set of cell-matrix adhesion genes that inversely follow claudin-5 levels are enriched in peripheral ECs.** Bar charts show the expression of the 12 cell-matrix adhesion genes identified in this study in ECs across organs in mice. Data from Munji *et al.*[26], plots were generated using *https://danemanlab.shinyapps.io/bbbtranscriptome/*. TPM: transcript per million. Note that all 12 of these genes are enriched in peripheral ECs and are relatively depleted from brain ECs in mice.

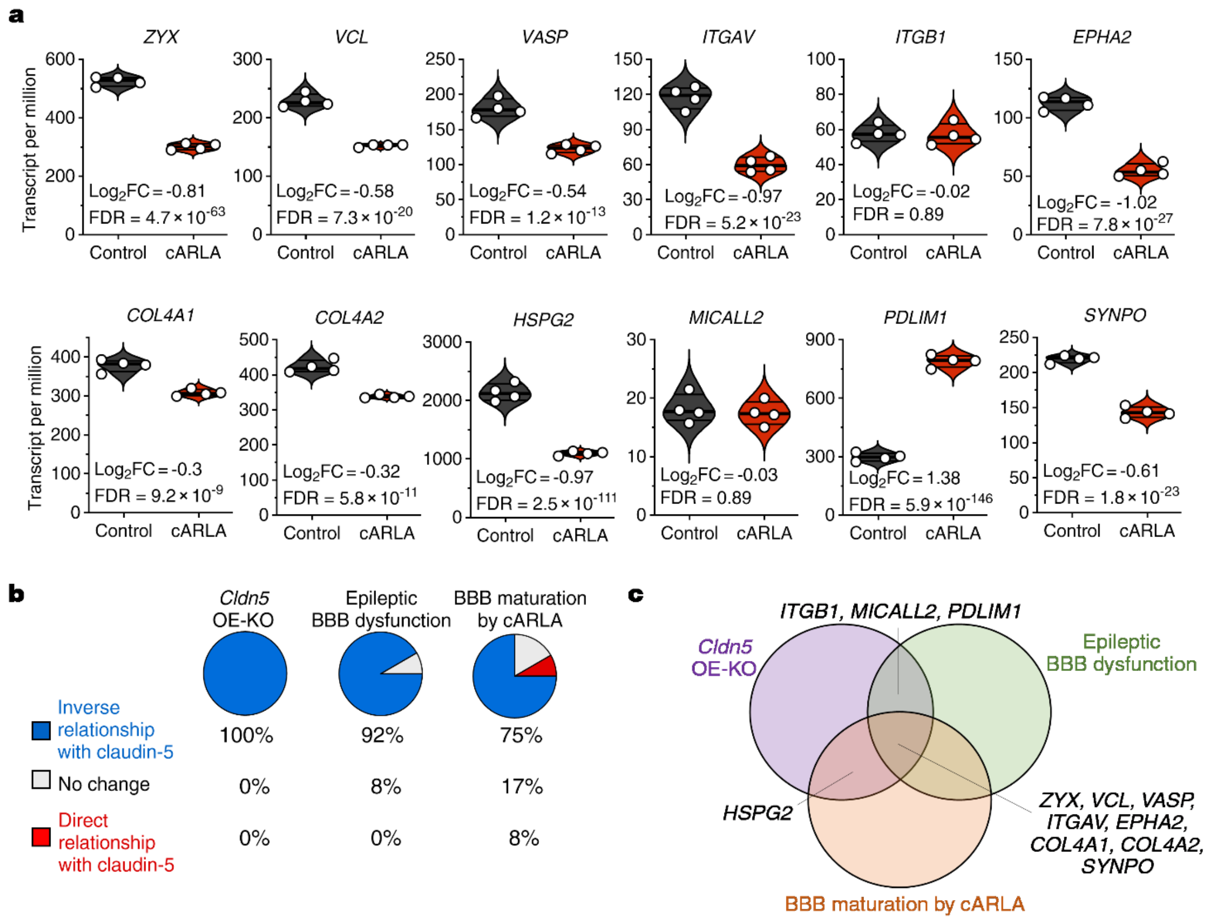


**Supplementary Fig. 9. Expression of the core set of cell-matrix adhesion genes that inversely follow claudin-5 levels across datasets. a)** Expression of genes in human stem cell-derived brain ECs with or without cARLA treatment. $Log_2FC$: $log_2$(fold change), FDR: adjusted *P*-value (Benjamini-Hochberg method). **b)** Pie charts showing the relationship of these 12 cell-matrix adhesion genes with claudin-5 levels across RNA-sequencing datasets used in this study. **c)** Venn diagram of shared and different gene regulation between the three datasets.

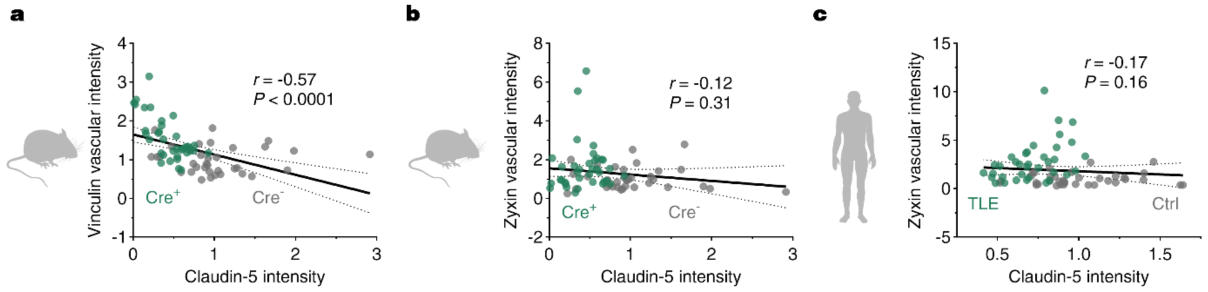


**Supplementary Fig. 10. Correlation of vessel-associated focal adhesion protein levels with claudin-5 levels as determined by confocal microscopy. a)** Correlation analysis of vessel-associated vinculin and claudin-5 levels in mouse brain sections, **b)** vessel-associated zyxin and claudin-5 in mouse brain sections and **c)** vessel-associated zyxin in human brain sections. For all three panels, Spearman correlation, two-tailed, n=72 datapoints from both control (grey) and TLE (green) groups. The line of best fit (continuous line) and 95% confidence intervals (dashed lines) from a simple linear regression are shown.

**Supplementary Table 1. Top upregulated pathways in mouse brain ECs in acute epilepsy.** Data reanalysed from Munji *et al.*[26], GO: Gene Ontology, BP: biological process, CC: cellular compartment, MF: molecular function, $P_{adj}$: adjusted *P*-value (FDR correction, Benjamini-Hochberg method). Gene ratio was calculated as the ratio of upregulated genes over the total genes in a given pathway. Pathways linked to cell-matrix adhesion and the actin cytoskeleton are highlighted in bold.

| Category | Name of GO term | Term ID | $-Log_{10}(P_{adj})$ | Gene ratio |
|---|---|---|---|---|
| **GO:CC** | **cell-substrate junction** | **GO:0030055** | **18,2154** | **0,2412** |
| **GO:CC** | **focal adhesion** | **GO:0005925** | **17,6479** | **0,2447** |
| GO:BP | extrinsic apoptotic signaling pathway | GO:0097191 | 15,8323 | 0,2137 |
| **GO:CC** | **actin filament bundle** | **GO:0032432** | **15,5758** | **0,3113** |
| **GO:BP** | **regulation of cell-substrate adhesion** | **GO:0010810** | **15,3149** | **0,2162** |
| **GO:MF** | **actin filament binding** | **GO:0051015** | **15,0430** | **0,2260** |
| GO:BP | endothelium development | GO:0003158 | 14,1427 | 0,2624 |
| GO:CC | ruffle | GO:0001726 | 14,1264 | 0,2253 |
| **GO:CC** | **actomyosin** | **GO:0042641** | **13,7747** | **0,2952** |
| **GO:CC** | **stress fiber** | **GO:0001725** | **13,0600** | **0,3021** |
| **GO:CC** | **contractile actin filament bundle** | **GO:0097517** | **13,0600** | **0,3021** |
| GO:BP | regulation of extrinsic apoptotic signaling pathway | GO:2001236 | 11,8175 | 0,2256 |
| GO:BP | endothelial cell differentiation | GO:0045446 | 11,4300 | 0,2627 |
| GO:BP | cellular response to vascular endothelial growth factor stimulus | GO:0035924 | 11,4018 | 0,3710 |
| GO:BP | response to hydrogen peroxide | GO:0042542 | 11,0989 | 0,2562 |
| GO:BP | positive regulation of supramolecular fiber organization | GO:1902905 | 10,9548 | 0,2077 |
| GO:BP | endothelial cell development | GO:0001885 | 10,9511 | 0,3380 |
| **GO:BP** | **positive regulation of cytoskeleton organization** | **GO:0051495** | **10,7445** | **0,2000** |
| **GO:BP** | **actin filament bundle assembly** | **GO:0051017** | **10,6836** | **0,2143** |

**Supplementary Table 2. Demographic details of temporal lobe epilepsy cases.** M: male, F: female, TLE: temporal lobe epilepsy, FA: focal aware, FIA: focal impaired aware, FBTCS: focal to bilateral tonic-clonic. Mean age at the time of surgery: 39.3.

| **Patient #** | **Sex** | **Age at surgery (years)** | **Type of epilepsy** | **Type of seizure** | **Seizure frequency** | **Duration of epilepsy (years)** | **Pathology** |
|---|---|---|---|---|---|---|---|
| 1 | M | 62 | TLE | 2 FIA | >1/month | 38 | Mild pyramidial neuronal loss |
| 2 | M | 35 | TLE | 1 FA, 2 FIA, 3 FBTCS | >1/month | 24 | Chaslin's subpial gliosis |
| 3 | F | 21 | TLE | 1 FA, 2 FIA, 3 FBTCS | >1/month | 9 | Chaslin's subpial gliosis |

**Supplementary Table 3. Demographic details of autopsy control cases.** Post-mortem hippocampal sections of non-diseased brains were provided by the Stanley Medical Research Institute. M: male, F: female. Mean age at the time of death: 35.3.

| Patient # | Sex | Age at death (years) | Post-mortem interval (hours) |
|---|---|---|---|
| 1 | M | 42 | 222 |
| 2 | M | 35 | 168 |
| 3 | F | 29 | 62 |

**Supplementary Table 4. List of antibodies and labeling probes used in this study.**

| Antibody / probe | Host | Use | Manufacturer, catalog number | Dilution |
|---|---|---|---|---|
| Alexa Fluor 488 anti-claudin-5 | mouse | brain tissue, confocal microscopy | Invitrogen, cat# 35-2588 | 1:200 |
| Anti-claudin-5 | mouse | dSTORM microscopy | Invitrogen, cat# 35-2500 | 1:200 |
| Anti-claudin-5 | rabbit | *in vitro*, confocal microscopy | Sigma-Aldrich, cat# SAB4502981 | 1:300 |
| Anti-claudin-5 | rabbit | brain tissue, confocal microscopy | Invitrogen, cat# 34-1600 | 1:200 |
| Anti-collagen IV | goat | brain tissue, confocal microscopy | Sigma-Aldrich, cat# AB769 | 1:100 |
| Anti-pFAK | rabbit | *in vitro*, confocal microscopy | Invitrogen, cat# PA5-78137 | 1:300 |
| Anti-vinculin | mouse | *in vitro*, confocal microscopy | Sigma-Aldrich, cat# V9264 | 1:300 |
| Anti-vinculin | rabbit | brain tissue, confocal microscopy | Invitrogen, cat# 700062 | 1:100 |
| Anti-vinculin | mouse | brain tissue, confocal microscopy | Sigma-Aldrich, cat# V9131 | 1:100 |
| Anti-zyxin | rabbit | *in vitro*, confocal microscopy | Sigma-Aldrich, cat# ABC1387 | 1:500 |
| Anti-zyxin | rabbit | brain tissue, confocal microscopy | Sigma-Aldrich, cat# ZRB1408 | 1:100 |
| Alexa Fluor 405 anti-goat | donkey | brain tissue, confocal microscopy | Invitrogen, cat# A-21235 | 1:500 |
| Alexa Fluor 488 phalloidin | - | *in vitro*, confocal microscopy | Invitrogen, cat# A12379 | 1:100 |
| Alexa Fluor 488 anti-rabbit | goat | *in vitro* + brain tissue, confocal microscopy | Invitrogen, cat# A-11008 | 1:500 |
| Alexa Fluor 555 anti-rabbit | goat | *in vitro* + brain tissue, confocal microscopy | Invitrogen, cat# A-21428 | 1:500 |
| Alexa Fluor 647 phalloidin | - | *in vitro*, confocal microscopy | Invitrogen, cat# A22287 | 1:100 |
| Alexa Fluor 647 anti-mouse | goat | dSTORM microscopy | Invitrogen, cat# cat# A-21235 | 1:1000 |
| Cy3 anti-mouse | goat | brain tissue, confocal microscopy | Invitrogen, cat# A10521 | 1:500 |
